# Neutrophils secrete exosome-associated DNA to resolve sterile acute inflammation

**DOI:** 10.1101/2024.04.21.590456

**Authors:** Subhash B. Arya, Samuel P. Collie, Yang Xu, Martin Fernandez, Jonathan Z. Sexton, Shyamal Mosalaganti, Pierre A. Coulombe, Carole A. Parent

## Abstract

Acute inflammation, characterized by a rapid influx of neutrophils, is a protective response that can lead to chronic inflammatory diseases when left unresolved. Secretion of LTB_4_-containing exosomes is required for effective neutrophil infiltration during inflammation. In this study, we show that neutrophils release nuclear DNA in a non-lytic, rapid, and repetitive manner, via a mechanism distinct from suicidal NET release and cell death. The packaging of nuclear DNA occurs in the lumen of nuclear envelope (NE)-derived multivesicular bodies (MVBs) that harbor the LTB_4_ synthesizing machinery and is mediated by the lamin B receptor (LBR) and chromatin decondensation. Disruption of secreted exosome-associated DNA (SEAD) in a model of sterile inflammation in mouse skin amplifies and prolongs the presence of neutrophils, impeding the onset of resolution. Together, these findings advance our understanding of neutrophil functions during inflammation and the physiological significance of NETs, with implications for novel treatments for inflammatory disorders.

## INTRODUCTION

A prompt and efficient response to tissue injury or infection lies at the core of the response to acute inflammation, which typically lasts a few hours^1,2^. Neutrophils, acting as first responders, infiltrate the tissue to clear harmful agents and damaged cells and secrete mediators essential for tissue repair^3,4^. Symptoms of acute inflammation, including redness, heat, swelling, and pain, usually subside after the initial phase of resolution led by neutrophils, and subsequently the involvement of macrophages and other immune cells^2,5^. In contrast, chronic inflammation, lasting weeks to years, often stems from unresolved acute inflammation, autoimmune responses, or persistent irritant exposure^6^. Temporal and/or functional dysregulation of neutrophil, macrophage, and lymphocyte infiltration is associated with several clinically significant chronic inflammatory conditions including rheumatoid arthritis, psoriasis, asthma, inflammatory bowel disease, chronic obstructive pulmonary disorder, systemic lupus erythematosus, atherosclerosis and cancer^6^. Many of these chronic conditions correlate with elevated levels of neutrophil extracellular traps (NETs) in both affected tissues and blood^7,8^. Classical, suicidal NETs are comprised of nuclear DNA embedded with toxic granular and cytoplasmic proteins and are released during necroptotic cell death triggered by either mitogen treatment, bacteria, or fungi^9,10^. In contrast, the non-lytic secretion of DNA during vital NET release has been reported to occur from phagocytosing neutrophils in human abscesses^11,12^. However, the precise mechanisms that regulate the non-lytic secretion of DNA and its role in inflammation remain unclear. While *in vitro* studies highlight the pro-inflammatory potential of NETs, the role of DNA secreted by viable neutrophils *in vivo* remains uncertain, with several studies pointing to their anti-inflammatory potential^13–16^.

The resolution of inflammation involves the clearance of pioneer neutrophils undergoing apoptosis, through efferocytosis by infiltrating macrophages^17^, in conjunction with the activation of peroxisome proliferator-activated receptors (PPARs) followed by upregulation of anti-inflammatory cytokines^18^. Furthermore, eicosanoids like lipoxins, leukotrienes, and prostaglandins contribute to the reverse transmigration of infiltrated neutrophils into the circulation, initiating the resolution response to ongoing inflammation^19,20^. Although the role of infiltrated neutrophils in initiating the resolution of inflammation is unclear, their role in promoting inflammation is well established, particularly in amplifying their recruitment range through the release of leukotriene B_4_ (LTB_4_) as they migrate toward injured or infected tissues^21,22^. LTB_4_ is synthesized from the cytosolic phospholipase A2α (cPLA2α)-dependent release of arachidonic acid from membranes through the serial action 5-lipoxygenase activating protein (FLAP; an endoplasmic reticulum/nuclear envelope-resident transmembrane protein), 5-lipoxygenase (5LO) and leukotriene A_4_ hydrolase (LTA_4_H)^23^. During neutrophil chemotaxis, LTB_4_-synthesizing enzymes and LTB_4_ itself are released within non-conventional exosomes generated from multivesicular bodies (MVBs) that originate from the nuclear envelope (NE) in a neutral sphingomyelinase (nSMase)-dependent manner^24,25^. LTB_4_ released from these exosomes acts both in an autocrine and paracrine fashion to amplify the recruitment of neutrophils to the infected/injured site^21,22^. Apart from its pro-inflammatory effects at low concentrations (∼10 nM), which ensues binding to a high-affinity GPCR, BLT1; higher concentrations (>100 nM) of LTB_4_ are required for the activation of a low-affinity GPCR, BLT2^26^. High concentrations of LTB_4_ are also required to activate the anti-inflammatory transcription factor, PPARα^27^, which participates in the resolution of both chronic and acute inflammation^28–30^. The precise mechanisms responsible for the spatiotemporal regulation of LTB_4_ levels within affected tissues and its role in influencing both neutrophil infiltration and PPARα-mediated resolution of inflammation remain unclear.

In this study, we aim to elucidate the significance of DNA secretion from migrating neutrophils in the context of inflammation and resolution. Our investigations identified a mechanism governing non-lytic DNA secretion from migrating neutrophils and the consequential impact of DNA secretion on neutrophil migration. Our findings reveal a role for the unique neutrophil NE composition in the secretion of both LTB_4_-containing exosomes and DNA. Moreover, we discovered that the secretion of exosome-associated DNA (SEADs) promotes the resolution of inflammation through the activation of the anti-inflammatory transcription factor PPARα.

## RESULTS

### Nuclear DNA is present in the lumen of NE-derived MVBs and is co-secreted with 5LO/FLAP-positive exosomes

Upon chemoattractant signaling, NE-derived 5LO/FLAP-positive MVBs originate from NE-buds that fuse with the plasma membrane (PM) to release NE-derived exosomes containing LTB_4_ and the LTB_4_ synthesis machinery^24,25^. The limiting membrane of NE-derived MVBs harbors lamin B receptor (LBR, an inner nuclear membrane transmembrane protein), while intraluminal vesicles (ILVs) lack LBR^24^. LBR plays a crucial role in cholesterol biosynthesis and the regulation of heterochromatin assembly^31–33^. In mature neutrophils of both human and mouse origin^34^, the levels of lamin A/C are low, but those of B-type lamins – particularly lamin B1 – are high^35^. The expression of LBR, which is essential for nuclear lobulation in neutrophils^36^, is also upregulated during neutrophil differentiation. LBR interacts with several proteins including the nucleosome core histone H3^37^. To gain further insight into the role of LBR within NE-derived MVBs, we immunostained human polymorphonuclear neutrophils (PMNs) actively migrating in an LTB_4_ gradient with anti-histone H3 antibodies. We observed strong histone H3 staining within LBR-positive NE-buds and cytosolic vesicles (**Fig. 1a-b**). Additionally, we found that structures positive for histone H3 are associated with 5LO-positive NE-buds and cytosolic vesicles corresponding to the size of NE-derived MVBs (**Fig. 1c-d**) ^24^. However, using 4x expansion microscopy we observe that the Hoechst and 5LO-positive ILVs present within the LBR-positive MVBs do not co-localize (**Fig. 1e**). Indeed, the low Pearson’s *R*-value between 5LO and Hoechst relative to LBR and Hoechst, measured using 3D-reconstruction of the expansion microscopy images (**Fig. 1f**), along with the luminal presence of chromatin-like structures in the MVBs of LTB_4_-stimulated human PMNs (**Fig. S1a-b**), suggest that DNA is associated with the LBR-positive limiting membrane of MVBs and is excluded from the interior of the ILVs.

**Figure 1.**
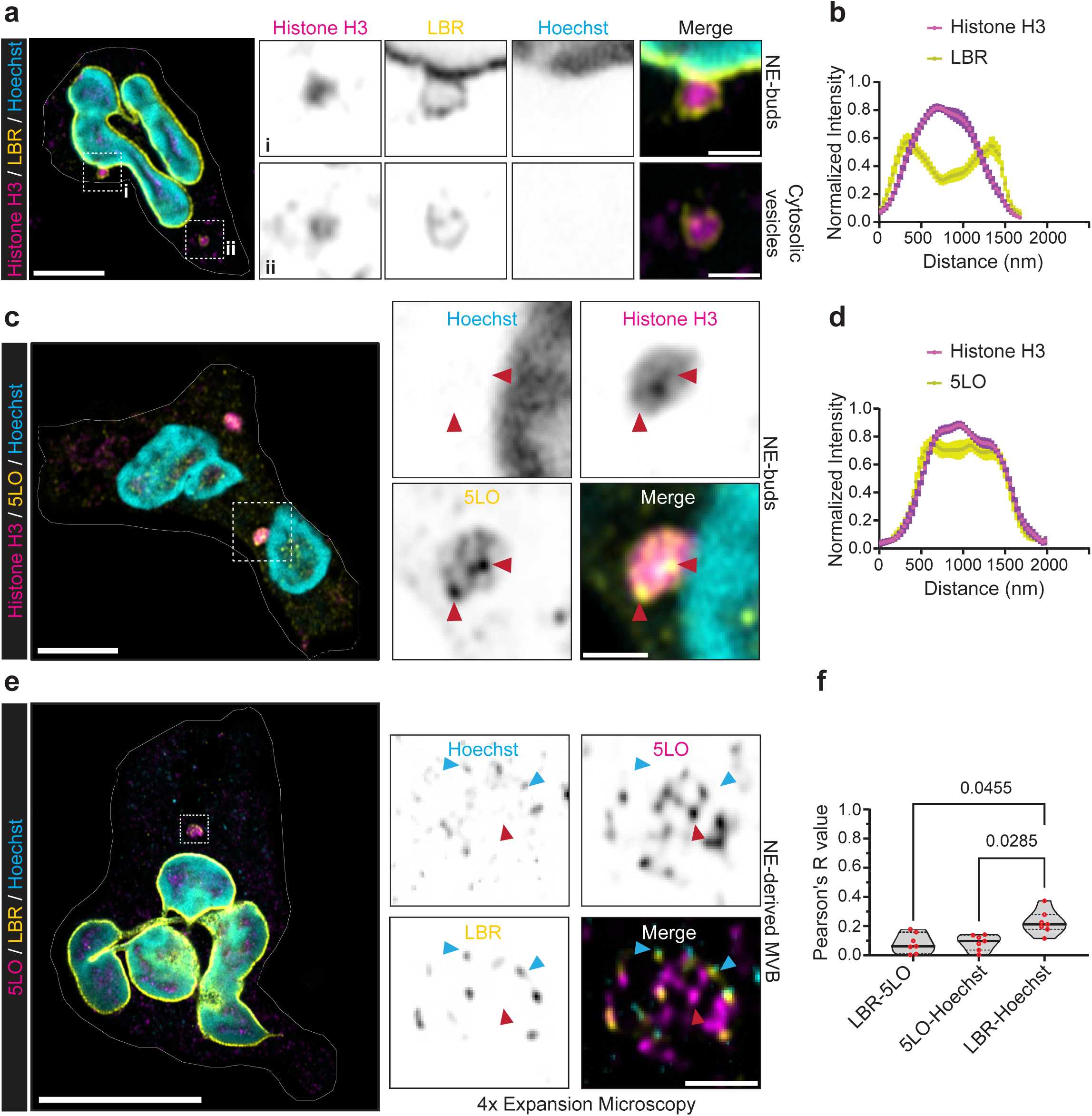
Nuclear DNA is packaged in the lumen of NE-derived MVBs in chemotaxing neutrophils. a. Representative Airyscan microscopy images of PMNs migrating towards LTB_4_, immunostained for LBR (yellow) and histone H3 (magenta), co-stained with Hoechst 33342 (cyan, nucleus). The scale is 5 µm and 1 µm in the inset. N=7. b. Histogram showing the normalized fluorescence intensity distribution of LBR and histone H3 across the diameter of LBR-positive NE-derived buds/vesicles. Data points from 26 different buds/vesicles obtained from 21 cells pooled from 7 independent experiments are plotted with the bold line showing the mean and the bars representing the s.e.m. c. Representative Airyscan microscopy images of PMNs migrating towards LTB_4_, immunostained for 5LO (yellow) and histone H3 (magenta), co-stained with Hoechst 33342 (cyan, nucleus). Red arrowheads point towards the region of intense 5LO staining. The scale is 5 µm and 1 µm in the inset. N=4. d. Histogram showing the normalized fluorescence intensity distribution of 5LO and histone H3 across the diameter of 5LO-positive buds/vesicles. Data points from 10 different buds/vesicles obtained from 8 cells pooled from 2 independent experiments are plotted with the bold line showing the mean and the bars representing the s.e.m. e. Four-fold expansion microscopy images of fixed PMNs chemotaxing towards 100 nM LTB_4_, immunostained for 5LO (magenta) and LBR (yellow), and co-stained with Hoechst 33342 (cyan). Blue and red arrowheads point towards Hoechst/LBR-positive structures and 5LO-positive punctae, respectively. The scale is 5 µm and 1 µm in the inset. N=3. f. Violin plot showing the distribution of Pearson’s *R*-value of either 5LO or LBR with Hoechst-positive structures and between LBR and 5LO structures, calculated using 3D-reconstructed multiple *z*-stack images of NE-derived buds/vesicles. The thick black line indicates the median. Red dots represent the data points collected from buds/vesicles of 7 different cells from 2 independent experiments.

To assess whether chromatin and NE-derived exosomes are co-secreted upon fusion of NE-MVBs with the plasma membrane, we employed cryogenic-corelative light electron microscopy (cryo-CLEM) to visualize the ultrastructure of NE-derived exosomes and co-secreted chromatin. We collected tilt-series at the periphery of plunge-frozen, LTB_4_-activated PMNs using the fluorescence from the membrane-impermeable DNA-binding dye SYTOXgreen as a guide (**Fig. 2a-b**). The reconstructed tomogram reveals the presence of vesicles ensconced within the secreted chromatin. Analysis of a single slice from the reconstructed tomogram shows the presence of SYTOXgreen-positive ‘bead-on-a-string chromatin-like’ structures associated with 130-200 nm diameter extracellular vesicles, proximal to intact and polarized PMNs (**Fig. 2c-d, Movie 1**). Furthermore, western blot analysis of exosomes purified from the supernatants of LTB_4_-activated PMNs shows the presence of histone H3 (chromatin marker) along with FLAP and 5LO (**Fig. 2e**), strongly supporting the co-secretion of chromatin with NE-derived exosomes. Notably, the histone H3 signal disappeared when exosomes were treated with DNase I (**Fig. 2e-f**), suggesting that the chromatin does not reside within the exosomes. Taken together, these results show that chromatin is present in the lumen of NE-derived MVBs, outside 5LO-positive ILVs, and is co-secreted with NE-derived exosomes containing the LTB_4_-synthesis machinery (**Fig. 2g**).

**Figure 2.**
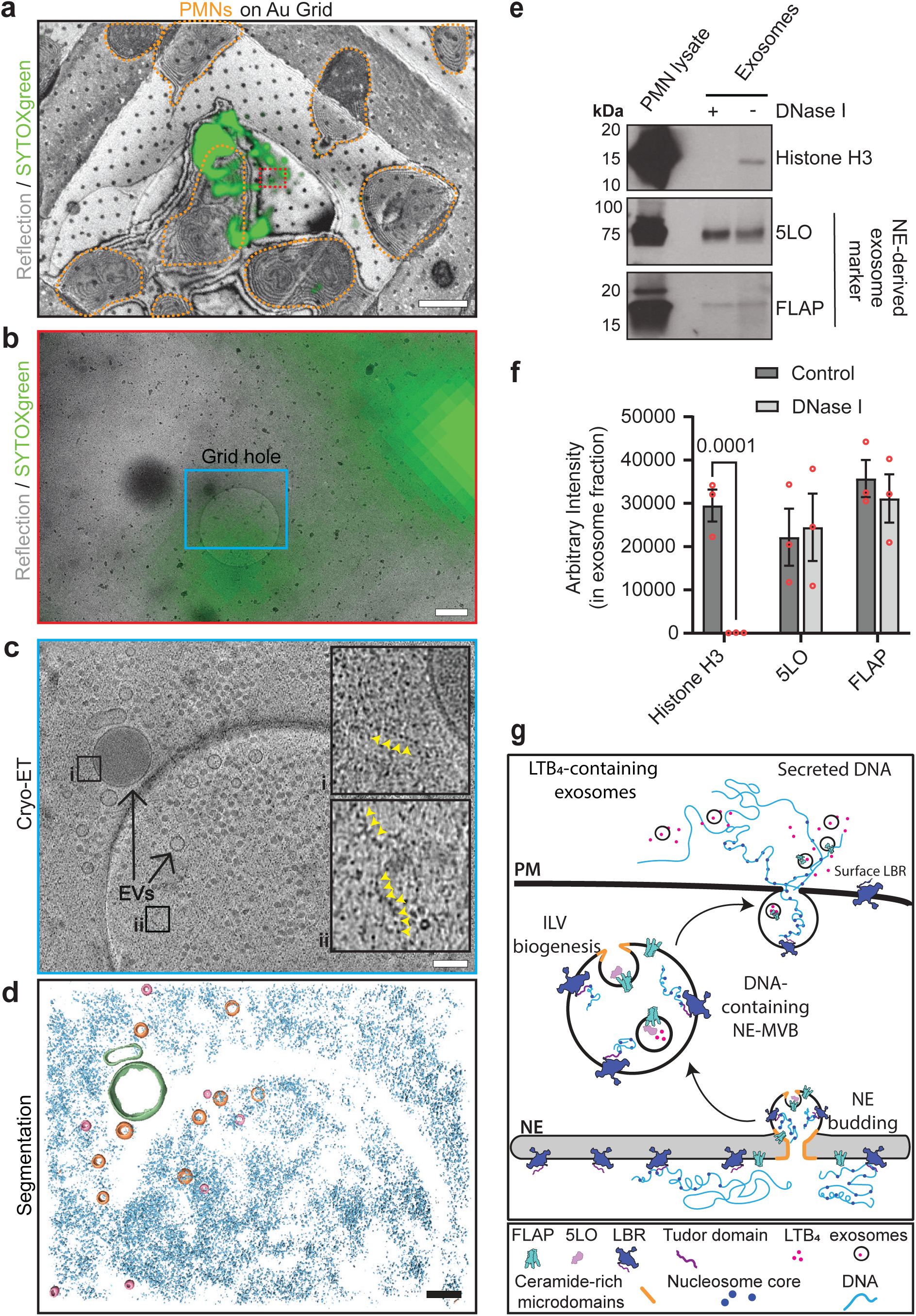
Spatial co-occurrence of nuclear DNA with NE-derived exosomes secreted from activated neutrophils. **a.** An overlay image of PMNs on fibrinogen-fibronectin coated Quantifoil grids stimulated with 100 nM LTB_4_ for 15 min in the presence of SYTOXgreen, plunge-frozen, and imaged on Leica Stellaris 5 cryo-confocal, showing the reflected light (gray) and extracellular DNA (green). Dashed yellow lines highlight PMNs on the grid square. The scale bar is 50 μm. **b.** A low-magnification TEM image (6500x) of the highlighted region (red box) in panel a, is overlaid with the SYTOX green fluorescence signal. The cyan box indicates the area where cryo-ET data was acquired. The scale bar is 500 nm. **c.** A slice through the tomogram of the region highlighted (cyan box) in panel b, with insets showing ‘bead-on-string’ like structures (yellow arrowheads). The scale bar is 100 nm. **d.** Segmentation of vesicles (<100 nm in orange, >100 nm in olive green) and the DNA (cyan) corresponding to the tomogram in panel c. The scale bar is 100 nm. See associated Movie 1. **e.** Representative western blot of exosomes obtained from fractions 4-9 of 5-40% density gradient centrifugation from PMNs stimulated with 100 nM LTB_4_ and treated with 50 µg/ml DNase I followed by immunoblotting for FLAP, 5LO, and histone H3. N=3. **f.** Bar graph showing the quantifications of the band intensity of FLAP, 5LO, and histone H3, in control and DNase I treated exosome samples. Data points (red dots) from 3 independent experiments were plotted as mean ± s.e.m. P values determined using two-way ANOVA are reported. **g.** Schematic illustration of DNA packaging and secretion during the nuclear budding and the fusion of NE-MVB with the PM. Note the change in the orientation of the LBR N-terminal Tudor domain as it moves from the inner nuclear membrane to the PM.

### Nuclear DNA is rapidly and repetitively secreted from the back of actively chemotaxing neutrophils

To determine whether chemotaxing neutrophils secrete DNA, we added SYTOXgreen in an under-agarose chemotaxis assay^38,39^ where Hoechst- and Cell Mask orange (plasma membrane)-stained PMNs are allowed to migrate toward LTB_4_. Remarkably, we observed that alive and actively migrating PMNs release DNA from their rear (**Fig. 3a, Fig. S2a, Movie 2**). Given the heterogeneity of neutrophils within individuals^40^ and across different donor populations^41,42^, we observed various phenotypes of DNA secretion. These phenotypes can be classified as (i) ‘DNA trails’ left by migrating neutrophils, (ii) ‘DNA blobs’ deposited by migrating neutrophils, and (iii) ‘attached-DNA blobs’ on the back of migrating neutrophils (**Movie 3**). Using CellProfiler-based object segmentation^43,44^, we classified neutrophils as positive for DNA secretion when they displayed the presence of DNA within 2 µm of the cell periphery for more than two minutes of migration. Using this approach, we determined that migrating cells exhibit no significant changes in several key nuclear morphological parameters describing lobulation (area extent, **Fig. S2c**) and surface complexity (form factor, **Fig. S3a**), either during or after DNA secretion. Migrating neutrophils maintained their PM integrity during DNA secretion, as supported by the absence of (i) SYTOXgreen staining in the cytoplasm of neutrophils actively secreting DNA in response to LTB_4_ (**Fig. S2a, c**), (ii) lysed cells quantified by cell segmentation analysis (**Fig. S2d**), and (iii) lactate dehydrogenase (LDH) activity in the supernatants of LTB_4_-treated PMNs (**Fig. S2e**); as well as the presence of consistent levels of intracellular calcium before, during, and after DNA secretion (**Fig. 3b**). Furthermore, the presence of nucleosome core histones in NE-buds and NE-derived cytoplasmic vesicles (**Fig. 1a, b**), along with the observed reduction in the nuclear Hoechst intensity measured after DNA secretion relative to the pre-secretion state (**Fig. 3c)**, show that the secreted DNA is of nuclear origin. To assess if the secretion of mitochondrial DNA through mitochondria damage also occurs^45,46^, we measured mitochondrial membrane potential in PMNs, using Mitotracker Red CMXrosamine^47^. We observed an increase in Mitotracker intensity during DNA secretion relative to before in PMNs migrating towards LTB_4_, suggesting that mitochondrial rupture is not associated with DNA secretion (**Fig. S2a, f**). Finally, we found that inhibiting nSMase activity significantly reduces the percentage of DNA-secreting PMNs (**Fig. 3d, Movie 4**), showing that DNA packaging within the NE-derived MVBs requires ceramide-rich membrane microdomain formation, as is the case for the biogenesis of LTB_4_-containing MVBs^24^.

**Figure 3.**
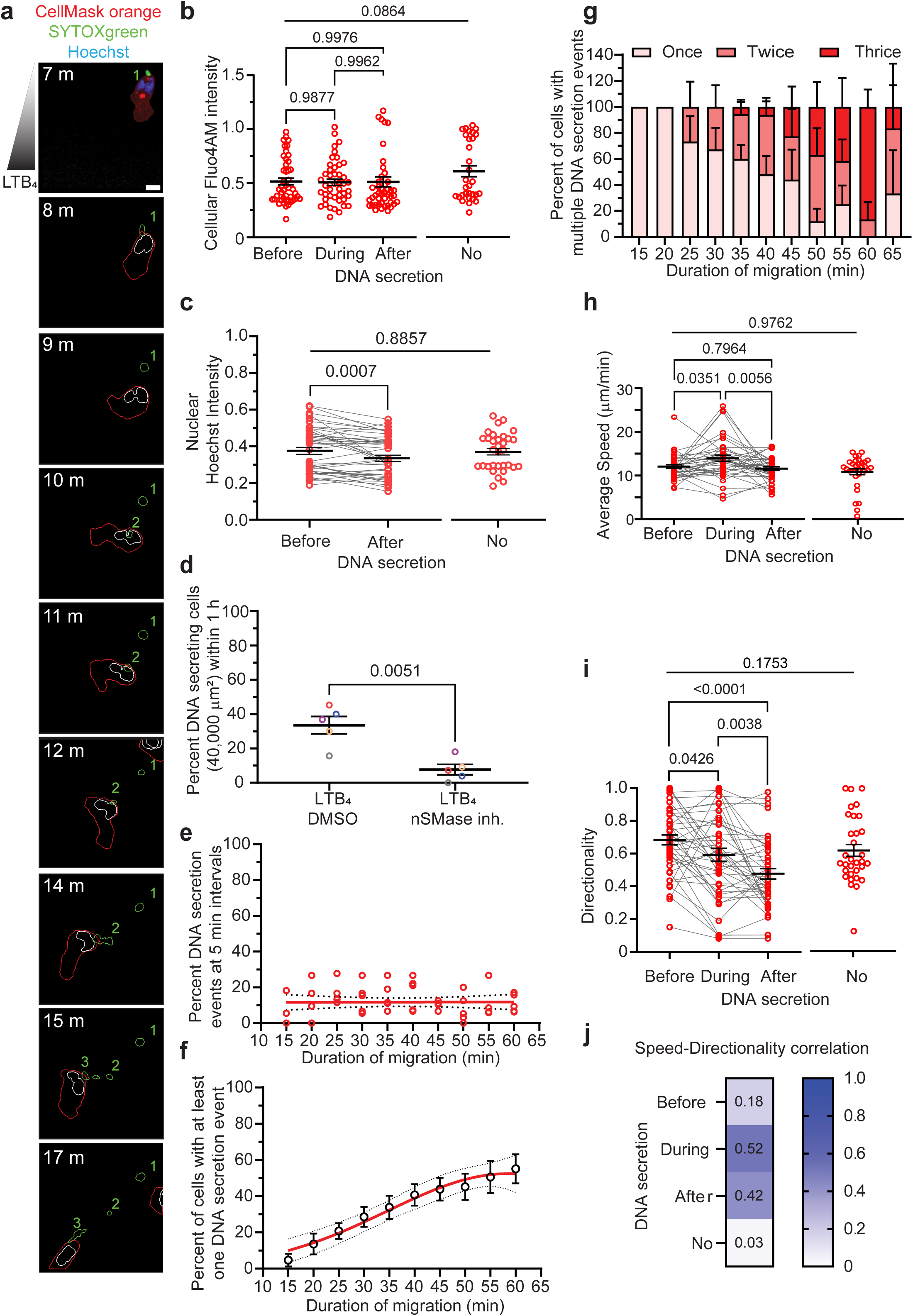
Fast and repetitive DNA secretion from the back of chemotaxing neutrophils. a. Representative Airyscan microscopy images of PMNs chemotaxing towards LTB_4_, showing the temporal dynamics of the PM (CellMaskOrange, red), and nucleus (Hoechst 33342, blue). Extracellular DNA is co-stained by SYTOXgreen (green). The depicted outlines for the cell (red), nucleus (white), and extracellular DNA (green) were generated using the CellProfiler image analysis tool. The number of times DNA is secreted is indicated in green on the image panels. The scale is 5 μm. See the associated Movie 2. N=5. b. Scatter dot plots showing the change in the cellular Fluo4-AM intensity (intracellular calcium) during DNA secretion in migrating PMNs. The red circles representing 45 datapoints for DNA-secreting cells and 31 data points for cells without DNA secretion are plotted as mean ± s.e.m., represented by the black lines. P values obtained using ordinary one-way ANOVA for DNA-secreting cells, and Mann-Whitney test for comparing before DNA secretion with no secretion conditions are plotted on the graph. N=5. c. Graph showing the fold change in average nuclear DNA (Hoechst) intensity in PMNs migrating towards LTB_4_ calculated for the duration of migration as mentioned on the x-axis. Datapoints (red circles) from 44 DNA-secreting cells are plotted as a before-after graph, and 32 cells with no DNA secretion are plotted as scatter dot plots. Thick black lines indicate mean ± s.e.m. P value calculated using ratio-paired t-test for before and after datapoints and Mann-Whitney test for before and no secretion data points are plotted. N=3. d. Scatter dot plot showing the changes in the percentage of DNA secreting PMNs within one hr, migrating towards LTB_4_ in the presence or absence of the nSMase inhibitor, GW4869, within an observation window of 40,000 μm^2^. P values calculated using paired t-tests are mentioned. Datapoints from the same experiments are color-coded similarly. See associated Movie 4. N=5. e. Scatter dot plot showing the percent of DNA secretion events of PMNs migrating towards LTB_4_ within 5-min intervals. Each circle represents data from one experiment. The solid red line represents the linear regression of the data, and the dotted black lines represent a 95% confidence interval. N=5. f. Graph showing the percentage of migrating PMNs with at least one DNA secretion event within an observation window of 40,000 μm^2^, calculated at 5 min intervals and plotted as mean ± s.e.m. The red line represents the non-linear regression determined using Gaussian least square fit and dotted gray lines show a 95% confidence interval. N=5. g. Bar graph showing the percent of PMNs with multiple DNA secretion events within 1 hr of migration in an area of 40,000 μm^2^, calculated every 5 min. Data from six independent experiments are plotted as mean ± s.e.m. h. Graph showing the changes in the average speed of migrating PMNs before, during, and after DNA secretion, and scatter dot plots of cells with no DNA secretion are plotted with thick black lines showing mean ± s.e.m. The red circles represent 42 datapoints for DNA-secreting cells and 32 datapoints for cells not secreting DNA. P value calculated using ordinary one-way ANOVA for before, during, and after data points and Mann-Whitney t-test for before and no secretion data points are plotted. N=5. i. Graph showing the changes in the directionality of migrating PMNs before, during, and after DNA secretion, and scatter dot plots of cells with no DNA secretion are plotted with thick black lines showing mean ± s.e.m. The red circles represent 46 data points for DNA-secreting cells and 32 datapoints for cells not secreting DNA. P value calculated using ordinary one-way ANOVA for before and after datapoints and Mann-Whitney test for before and no secretion data points are plotted. N=5. j. Image showing the speed-directionality correlation at different phases of PMN chemotaxis.

This vital DNA secretion response sharply contrasts with the behavior of PMNs undergoing PMA-induced suicidal NET release, which involves nuclear de-lobulation, nuclear swelling, cell rupture, and subsequent release of large amounts of nuclear DNA^48^ (**Fig. S2b-d, Movie 5**). We also found that, unlike PMA-induced suicidal NET release^49,50^, the secretion of DNA from actively chemotaxing PMNs does not require PAD4 or NOX2 activity (**Fig. S2g, Movie 6**). Similarly, unlike the dependence of PMA-induced NET release on RIPK3 and Gasdermin D signaling pathways^51,52^, pharmacological perturbation of the necroptosis mediator RIPK3 does not alter the DNA-secreting potential of PMNs migrating towards LTB_4_ (**Fig. S2h**). However, and not surprisingly, since LTB_4_ signaling exerts an anti-apoptotic effect on neutrophils^53,54^, we found that inhibiting caspase 3 activity increases the percentage of DNA-secreting neutrophils migrating toward LTB_4_(**Fig. S2h, Movie 7**)). Intriguingly, we did observe that inhibition of ferroptosis tends to decrease DNA secretion (**Fig. S2h, Movie 7**), although both the necroptosis and ferroptosis inhibitors had observable defects in neutrophil migration.

Further characterization of the kinetics of DNA release revealed that, at any given moment within an hour of migration towards LTB_4_, the percentage of chemotaxing PMNs secreting DNA remains constant (**Fig. 3e**). By comparison, the cumulative percentage of PMNs that secreted DNA at least once within an imaging window of 40,000 μm^2^ increases proportionally with time, until reaching saturation to ∼50% (**Fig. 3f**). We also found that among DNA-secreting cells, the percentage of cells that secrete DNA twice or more increases with the duration of migration (**Fig. 3g**) and that DNA secretion events are accompanied by an increase in average cell speed, followed by a return to the pre-secretion speed matching that of cells leaving no DNA tracks (**Fig. 3h**). This increase in speed is not accompanied by changes in cell polarization (**Fig. S3b**). However, a subset of neutrophils did exhibit an increase in directionality during DNA secretion relative to before and after secretion (**Fig. 3i**). That said, we measured a higher correlation between speed and directionality in DNA-secreting neutrophils compared to neutrophils with no DNA secretion (**Fig. 3j**). Together, these findings show that chemotaxing PMNs secrete nuclear DNA in a non-lytic, rapid, and repetitive manner that is dependent on nSMase-mediated NE budding.

### LBR is crucial for the packaging and secretion of DNA but not for the biogenesis and secretion of NE-derived exosomes

Human promyelocytic leukemia-derived cells, HL60, can be readily differentiated into neutrophil-like cells when cultured in the presence of DMSO^38,55,56^. Unlike PMNs, differentiated HL60 (dHL60) cells lack nuclear lobulation, express high levels of lamin A, and exhibit a predominantly cytoplasmic LBR localization (**Fig. S4a-e**). To generate an HL60-derived variant that more closely resembles PMNs, we used CRISPR-cas9 genome editing to create an *LMNA* knockout (KO) HL60 cell line. Relative to scrambled (SCR) dHL60 cells, *LMNA* KO dHL60 cells show an improved NE distribution of LBR, enhanced nuclear lobulation and heterochromatin reorganization, much like PMNs (**Fig. S4a-f**). Proteomic analysis shows that compared to SCR dHL60 cells, *LMNA* KO dHL60 cells exhibit higher steady-state levels of key proteins associated with neutrophil activation, degranulation, and exocytosis, and lower steady-state levels of proteins related to chromatin organization, DNA replication, and RNA translation (**Fig. S5a-b**). As expected, relative to SCR controls the *LMNA* KO dHL60 cells exhibit more robust and faster migration towards N-formylMethionine-Leucyl-Phenylalanine (fMLF), a primary chemoattractant (**Fig. S5c-d**), with no change in either directionality of migration (**Fig. S5e**) or cell polarization (**Fig. S5f**).

We next generated *LMNA/LBR* KO cells and found that they display spherical/ovoid nuclei, similar to the previously reported dHL60 *LBR* KO cells^57^; in addition, we observed a decrease in heterochromatin segregation in *LMNA/LBR* KO cells relative to *LMNA* KO dHL60 cells (**Fig. S4a-f**). Furthermore, *LMNA/LBR* KO cells show a decrease in the number of cells migrating towards fMLF and in the median speed relative to *LMNA* KO cells, although no change in directionality or polarity was observed (**Fig. S5c-f**). In this context, the nuclear morphology of *LMNA/LBR* KO dHL60 cells nicely mimics the one observed in neutrophils from individuals with Pelger Huet anomaly (PHA)^58^, an inherited condition harboring LBR mutations that results in hypo-segmented neutrophil nuclei and increased susceptibility to specific infections^59,60^. We, therefore, used the *LMNA* KO and *LMNA/LBR* KO cells to characterize the role of LBR in the biogenesis of NE-derived exosomes and DNA release.

We first employed density-gradient ultracentrifugation to compare the composition of MVBs in *LMNA* KO and *LMNA/LBR* KO dHL60 cells treated with LTB_4_. We found that in *LMNA* KO cells, the NE-derived MVB markers, FLAP and LBR, co-fractionate with CD63, a marker of conventional PM-derived MVBs (**Fig. 4a-b**). Moreover, we found that while the chromatin marker histone H3 is present in the MVB fraction of *LMNA* KO dHL60 cells, it is significantly depleted in the *LMNA/LBR* KO mutants (**Fig. 4a-b**). Western blotting of exosomes isolated from LTB_4_-treated *LMNA* KO and *LMNA/LBR* KO dHL60 cells confirmed that LBR is critical for the secretion of chromatin but not NE-derived exosomes since we observe a dramatic loss of histone H3 signal in exosomes isolated from *LMNA/LBR* KO dHL60 cells (**Fig. 4c-d**). Notably, we found that LBR is absent in exosomes, further confirming its role in tethering chromatin to the limiting membrane of NE-derived MVBs (**Fig. 4c-d** and **Fig. 2g**). Given that the N-terminal Tudor domain of LBR mediates its binding to histone H3/H4^32^, we next subjected dHL60 cells to flow cytometry using an antibody directed against the first 200 amino acids of LBR to assess surface exposure of this domain. We measured a significant increase in both the percentage of cells positive for surface LBR (**Fig. 4e, f**) and in the mean fluorescence intensity (MFI) of surface LBR (**Fig. 4g**) in LTB_4_-activated SCR dHL60 cells, compared to DMSO-treated SCR dHL60 cells. *LBR* KO dHL60 cells served as a negative control for anti-LBR antibody staining. Finally, as anticipated, we observed that the percentage of cells secreting DNA is significantly reduced in *LMNA/LBR* KO dHL60 cells migrating towards LTB_4_, compared to *LMNA* KO cells (**Fig. 4h, Movie 8**). Together, these results highlight the crucial role of LBR in the packaging of histone H3-positive chromatin in NE-derived MVBs and their subsequent secretion, with no apparent involvement in the biogenesis and secretion of NE-derived exosomes.

**Figure 4.**
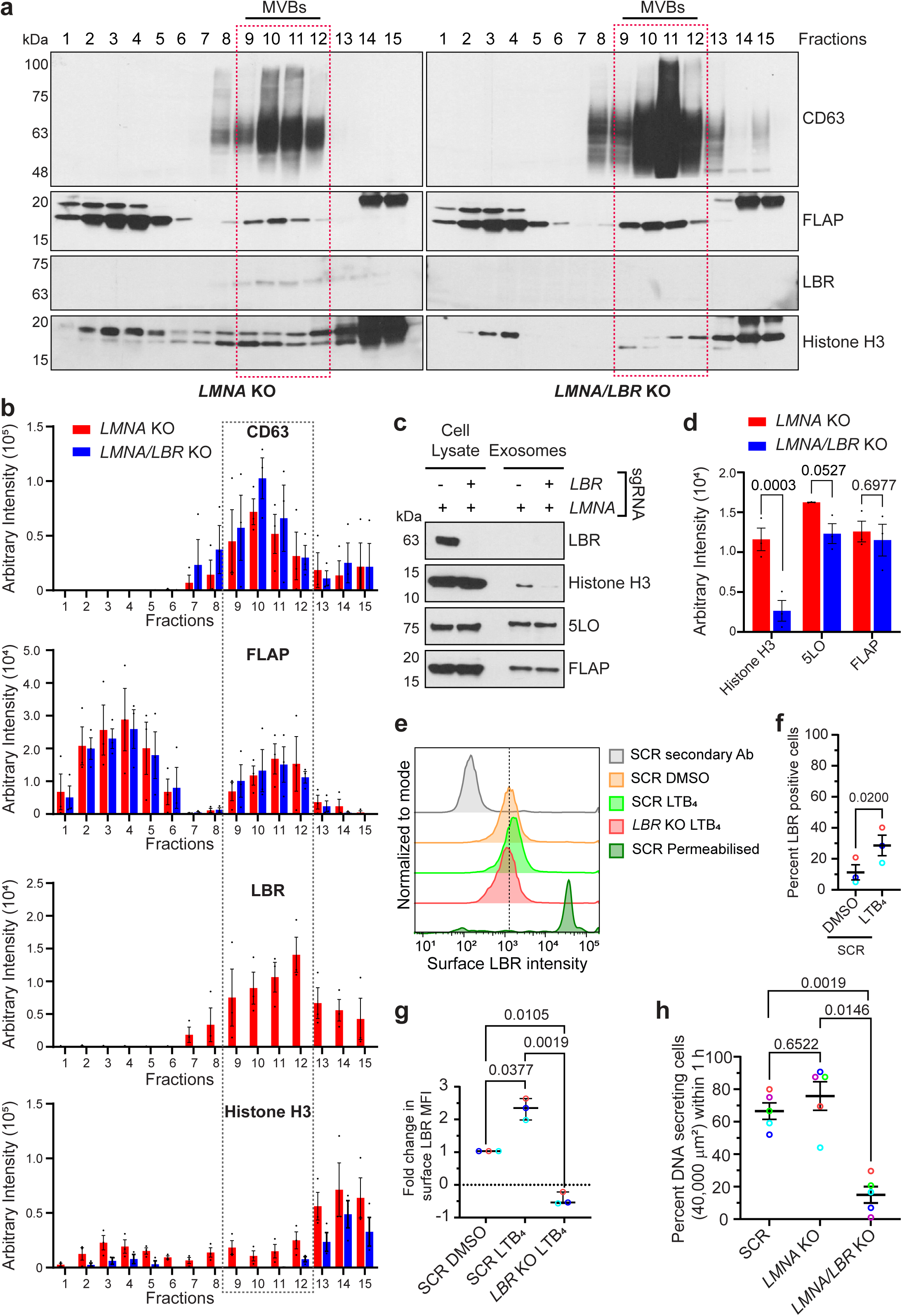
LBR is required for the packaging and secretion of DNA, but not exosomes, through NE-derived MVBs. a. Representative western blot images showing the levels of CD63, FLAP, LBR, and histone H3 expression in different subcellular fractions obtained by density-gradient ultracentrifugation of cytoplasmic contents of *LMNA* and *LMNA/LBR* KO dHL60 cells treated with LTB_4_ (100 nM) for 30 min. The red box highlights the MVB fractions. Molecular weights - in kiloDaltons (kDa) - are located on the left side of the panels. N=3. b. Bar graphs showing the intensity levels of proteins quantified using western blot images as shown in panel a. Data is plotted as mean ± s.e.m. N=3. c. Representative western blot images showing the levels of LBR, histone H3, 5LO, and FLAP intensities in the cell lysates and purified exosomes obtained from *LMNA* KO and *LMNA/LBR* KO dHL60 cells treated with LTB_4_ (100 nM) for 30 min. Molecular weights in kDa are located on the left side of the panels. N=3. d. Bar graphs showing changes in the intensity levels of proteins in exosome fractions as shown in panel c. Data is plotted as mean ± s.e.m. and P values calculated using two-way ANOVA analysis are shown. N=3. e. Histogram showing the intensity of surface LBR staining in SCR dHL60 cells treated with either DMSO or 100 nM LTB_4_ and immunostained for LBR, under non-permeabilizing conditions. Fixed, TritonX-100 permeabilized SCR cells were used as a positive control to stain intracellular LBR, and *LBR* KO cells were used as a negative control. f. Scatter dot plot showing the percentage of LBR (surface)-positive SCR dHL60 cells post LTB_4_ activation. Datapoints from the same experiments are similarly color-coded and are plotted as mean ± s.e.m. and P values calculated using paired t-tests are shown. N=3. g. Scatter dot plot showing the relative change in the surface LBR intensity within indicates conditions (x-axis). Datapoints from the same experiments are similarly color-coded and are plotted as mean ± s.e.m. and P values calculated using RM one-way ANOVA are shown. N=3. h. Scatter dot plot showing the percent of DNA-secreting cells within SCR, *LMNA* KO, and *LMNA/LBR* KO dHL60 cells migrating towards LTB_4_, within an observation window of 40,000 μm^2^ for 1 hr. Datapoints from the same experiments are similarly color-coded and are plotted as mean ± s.e.m., and P values obtained using paired t-tests are shown. N=5. See associated Movie 8.

### LTB_4_-induced histone acetylation promotes DNA secretion

Acetylation of histone H3 at lysine 27 (H3K27ac) has been reported as one of the most notable and rapidly elicited changes during activation-induced chromatin unwinding events in zebrafish neutrophils^61^ (**Fig. 5a**). Increases in histone acetylation have otherwise been reported to promote both NOX2-dependent and independent NET release^62^. Using western blot analysis of acid-extracted histones, we found that LTB_4_ induces an increase in the levels of H3K27ac relative to DMSO-treated PMNs (**Fig. 5b-c**). Pretreatment with anacardic acid and trichostatin A, individually, employed here as controls to respectively inhibit histone acetylase (HAT) and histone deacetylase (HDAC)^63^, altered H3K27Ac levels without changing the levels of citrullinated histone H3 during PMN activation (**Fig. 5b-c**). We also observed that H3K27ac co-localizes with 5LO-positive cytosolic vesicles in chemotaxing PMNs (**Fig. 5d**). Furthermore, object segmentation analysis of LBR-positive NE-derived buds and cytosolic vesicles revealed an increase in the intensity of histone H3 within these structures in HDAC inhibited PMNs migrating towards LTB_4_, compared to DMSO-treated PMNs (**Fig. 5e**). Conversely, we found that HAT inhibition decreases the abundance of histone H3 within NE-derived vesicles (**Fig. 5e**). By contrast, no change occurs in the size of the cytosolic vesicles occurs upon perturbation of either HAT or HDAC activity (**Fig. 5f**). Furthermore, PMNs pretreated with the HDAC inhibitor exhibit a significant increase in the percentage of DNA-secreting cells migrating towards LTB_4_, with a proportional decrease upon HAT inhibition (**Fig. 5g, Movie 9**). Together, these findings suggest that DNA packaging within NE-derived vesicles is mediated through LTB_4_-induced histone acetylation in neutrophils.

**Figure 5.**
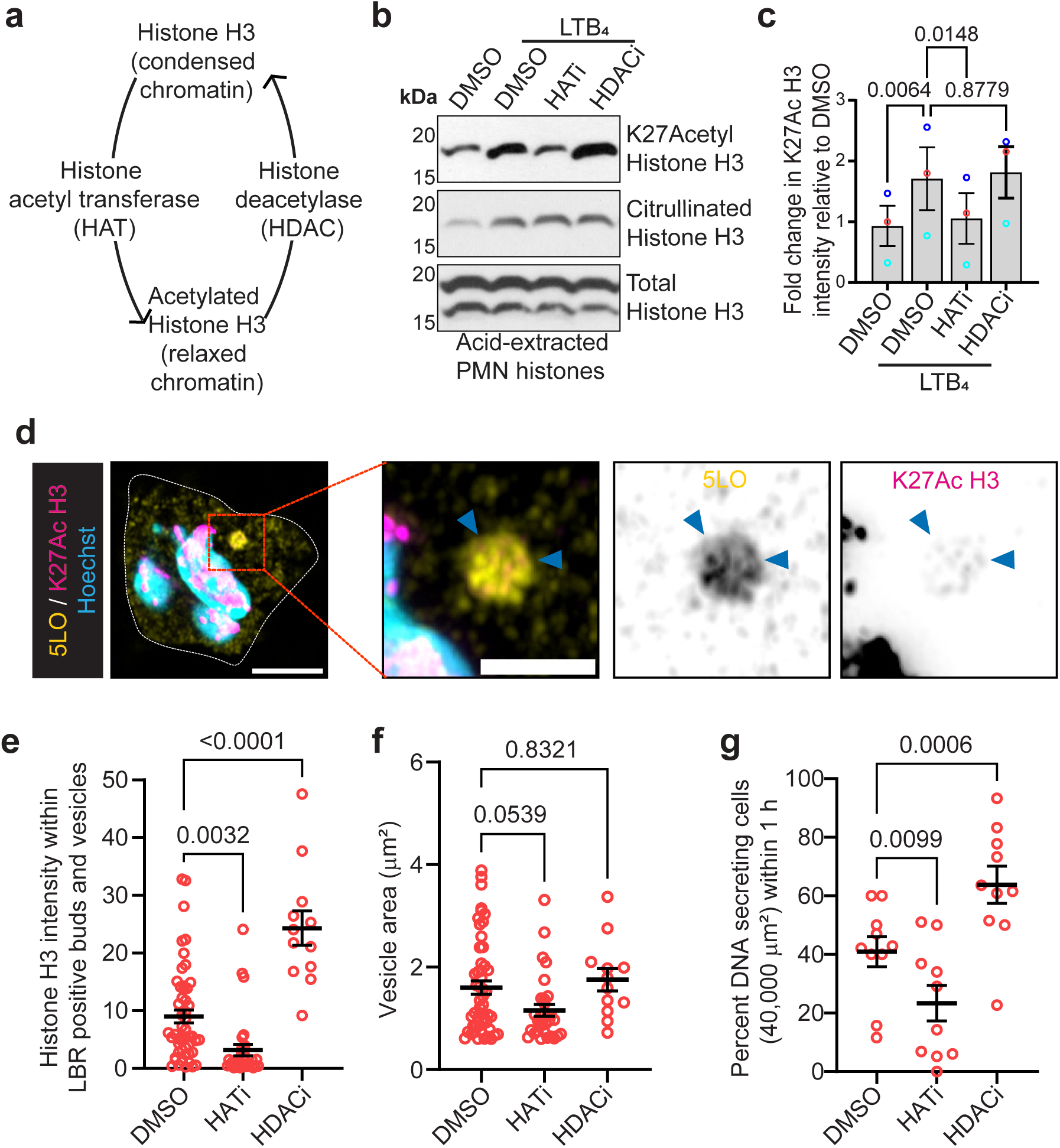
LTB_4_-induced histone acetylation mediates DNA secretion. a. Schematic depicting the effect of histone acetylation modifiers on chromatin architecture. b. Representative western blot images showing the levels of histone H3 K27acetyl, total histone H3, and citrullinated histone H3 signal intensity, in acid-extracted histone fractions, obtained from PMNs treated with either DMSO or 100 nM LTB_4_, for 5 min in the presence or absence of HAT and HDAC inhibitors. N=3. c. Bar graph showing the fold change in the levels of histone H3 K27acetyl normalized to total Histone H3 levels in different samples, normalized with that of DMSO from a single experiment. Datapoints from the same experiments are similarly color-coded and are plotted as mean ± s.e.m., and P values calculated using RM one-way ANOVA are shown. N=3. d. Representative Airyscan microscopy images of PMNs migrating towards LTB_4_, fixed and immunostained for 5LO (yellow), histone H3 K27 acetyl (magenta), and Hoechst 33342 (cyan, nucleus). Blue arrowheads in the zoomed insets point towards H3K27Ac enrichment within NE-derived MVB. The scale is 5 μm, in the inset, it is 1 μm. N=3. e. Scatter dot plots showing total histone H3 intensity within LBR positive buds and vesicles in PMNs migrating towards LTB_4_ under the conditions on the x-axis, within 1 hr. Data quantified using segmentation and analysis platform CellProfiler are plotted as mean ± s.e.m. of 52, 31, and 12 data points (red circles) belonging to DMSO, HAT inhibitor, and HDAC inhibitor-treated samples, respectively, pooled from 5 independent experiments. P values calculated using ordinary one-way ANOVA are shown. f. Scatter dot plots showing the area of LBR-positive buds/vesicles from the same images quantified and plotted similarly to panel e. g. Scatter dot plot showing the percent of DNA-secreting PMNs migrating towards LTB_4_, within an observation window of 40,000 μm^2^, for 1 hr, in the presence or absence of either HAT or HDAC inhibitors. Datapoints (red circles) are plotted as mean ± s.e.m. and P values obtained using mixed-effect analysis are shown. N=10. Also, see Movie 9.

### Secreted DNA promotes neutrophil resolution in inflamed mouse skin

To investigate the impact of secreted DNA on neutrophil chemotaxis, we compared the ability of PMNs to migrate towards LTB_4_ *ex vivo* in the presence or absence of DNase I. Cell tracking analysis revealed that DNase I-mediated digestion of secreted DNA gives rise to a robust increase in the number of migrating neutrophils, as well as a decrease in their directionality, although the average migration speed remains unchanged (**Fig. 6a-d, Movie 10**). These results are similar to what has been reported with neutrophils treated with inhibitors of LTB_4_ synthesis^22,25^. Since chemotaxing neutrophils co-secrete LTB_4_-containing exosomes and DNA, we envision that DNase I treatment results in the dispersion of LTB_4_-containing exosomes, thereby affecting autocrine and paracrine signaling. To test this, we inhibited LTB_4_ biogenesis using MK886, a FLAP inhibitor^64,65^. As expected, we found a decrease in the directionality of PMN chemotaxis upon MK886 pre-treatment, an effect that phenocopies DNase I treatment (**Fig. 6a-d**). Moreover, inhibiting LTB_4_ biogenesis in PMNs migrating in the presence of DNase I did not further affect the decline in chemotaxis directionality (**Fig. 6a-d**). Together, these results suggest that the secretion of DNA from migrating neutrophils regulates the directionality of PMNs in an LTB_4_-dependent manner. We envision that this could be through the clustering of LTB_4_-containing exosomes on the released DNA, thereby facilitating LTB_4_-mediated autocrine and paracrine signaling.

**Figure 6.**
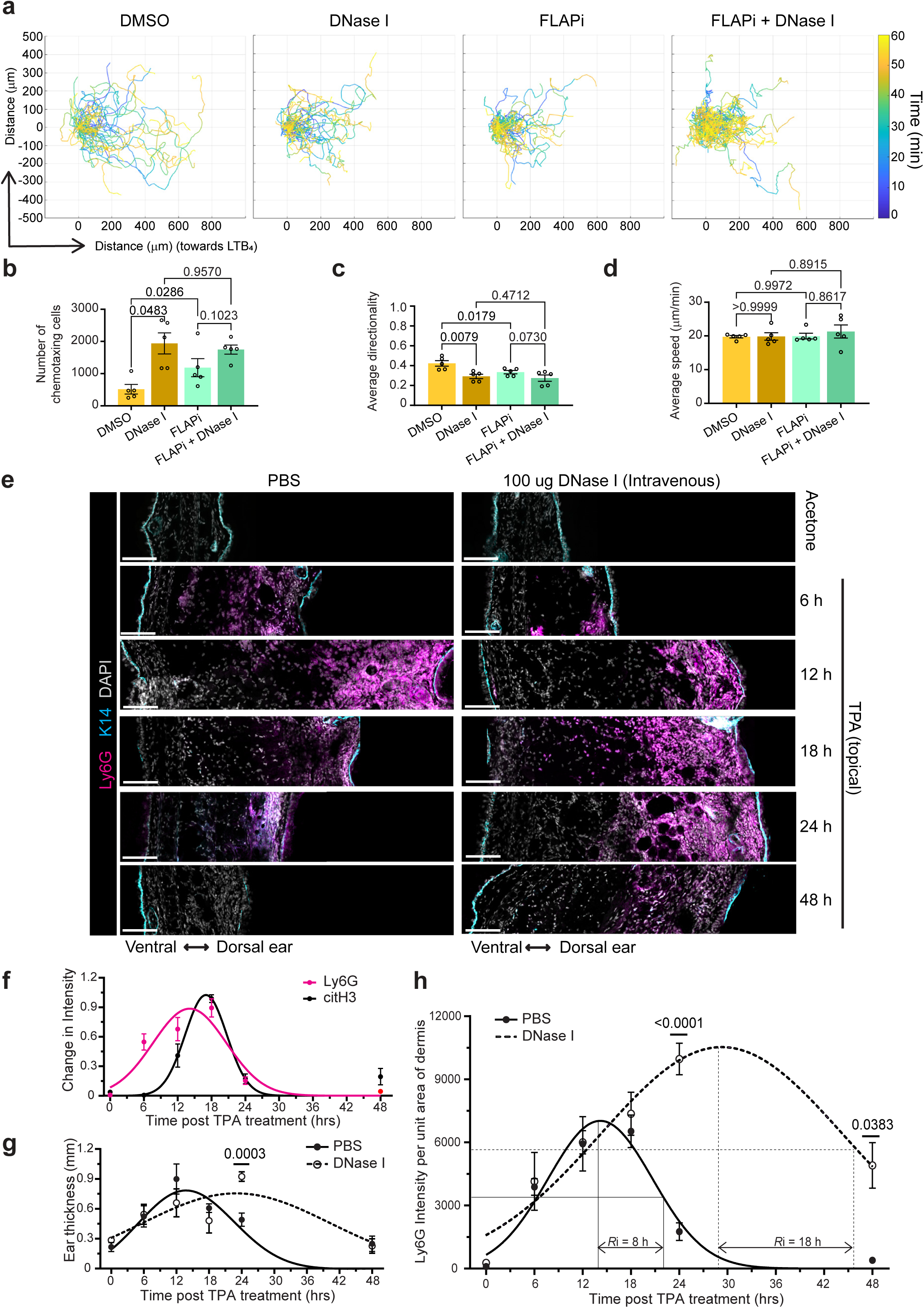
Disruption of secreted DNA increases neutrophil chemotaxis *in vitro* and infiltration to inflamed sites *in vivo*. **a.** Representative cell tracks of PMNs pretreated with either DMSO (vehicle) or MK886 (1 μM), migrating towards LTB_4_ (x-axis) in the presence or absence of DNase I (10 U/ml) for an hour. Graphs display the migration of 100 randomly selected tracks from each condition. The color-coded time scale is shown on the right. N=5. See associated Movie 10. **b-d.** Bar graphs showing the number of PMNs migrated towards LTB_4_ within 1 hr **(b**), average directionality (**c**), and average speed (**d**) in the presence or absence of DNase I. Datapoints (black circles) are plotted as mean ± s.e.m. and P values calculated using RM one-way ANOVA are shown. N=5. **e.** Representative Airyscan microscopy images of acetone/TPA-treated mice ear cryosections (20-micron) fixed and stained for Ly6G (magenta, neutrophils), K14 (cyan, epidermis), and DAPI (gray, nuclei) showing the ‘sum of slices’ projection of the optical slices captured every 600 nm throughout the tissue thickness. The scale is 100 μm. N=3. **f.** Graph showing the temporal kinetics of extracellular DNA and infiltrated neutrophils in the dermis of TPA-treated inflamed ears. Datapoints (magenta/black dots) are plotted as mean ± s.e.m. Thick colored lines (magenta/black) demonstrate the non-linear Gaussian-curve fit of the data. **g-h.** Graphs showing the temporal changes in the thickness of TPA-treated ears (**g**) and neutrophil infiltration in the ear dermis (**h**) from mice injected intravenously with either PBS or 100 μg DNase I, 1 h prior to TPA treatment. Thin dotted lines highlight the approach used to calculate the resolution interval (*R*i) in panel h. Datapoints are plotted as mean ± s.e.m. and P values calculated using two-way ANOVA analysis are shown. Thick solid/dotted lines (black) demonstrate the non-linear Gaussian-curve fit of the data. N=3.

To assess the effect of secreted DNA on the kinetics of neutrophil infiltration *in vivo*, we took advantage of a well-established model of acute skin inflammation in mice, induced by topical application of a phorbol ester, tetradecanoylphorbol-13-acetate (TPA)^66,67^. The primary targets of topically applied TPA are the suprabasal differentiating keratinocytes of the epidermis^67^. DNase I or PBS control, was injected in the tail vein one hour prior to topical TPA treatment on the dorsal ear skin. In the PBS-injected control mice, topical TPA treatment resulted in a robust but transient inflammation response that included a significant thickening of the skin tissue, particularly the dermis (connective tissue stroma), and a concomitant dermis-restricted neutrophil infiltration (assessed using Ly6G staining). We found that neutrophil infiltration peaks at 12 h and returns to basal levels by 48 h (**Fig. 6e-g**) - kinetics that align with previous studies^67–69^. The absence of Ly6G staining on the ventral (untreated) side of the ear indicates that neutrophil infiltration occurs in the tissue proximal to the TPA-treated area. Using an antibody against citrullinated histone H3^70,71^, we also readily detected^58–60^ extracellular DNA by 12 h, with a peak at ∼18 h and return to basal levels by 48 h after topical TPA treatment in mice ear (**Fig. 6f and S6a**). As expected, no signal for citrullinated histone H3 is detected in the dermis of DNase I-injected mice after TPA treatment (**Fig. S6a**). Furthermore, we observed that in DNase I-injected mice, TPA treatment elicits a delayed but sustained inflammatory response, as measured by ear tissue thickness (**Fig. 6e, g**), and prolonged neutrophil infiltration in the ear dermis peaking at ∼30 h post-TPA treatment (**Fig. 6e, h**). While infiltrated neutrophils are not detected in the dermis of TPA-treated ears after 48 h in PBS-injected mice, as much as ∼50% of infiltrated neutrophils persist at this time point in DNase I-injected mice, resulting in more than 2-fold increase in the resolution interval (*R*i)^72^ (**Fig. 6h**). Finally, we also observed that the DNase I treatment inhibits the infiltration of monocytes as assessed via F4/80 staining (**Fig. S6b**) - a process known to coincide with the initiation of neutrophil resolution^73,74^. Together, these results establish a role for secreted extracellular DNA in promoting the resolution of infiltrated neutrophils from the site of an acute sterile inflammation.

We next sought to further determine the nature of secreted DNA by visualizing the spatial co-occurrence of NE-derived exosomes and secreted DNA in inflamed mouse ears. Considering that FLAP is a transmembrane protein^64^ and an integral component of NE-derived exosomes^24,25^, we used a monoclonal antibody directed against FLAP to detect these exosomes in inflamed ears. We also used an anti-double-stranded DNA (dsDNA) antibody to assist the tracking of secreted extracellular DNA. To visualize the spatial co-occurrence of NE-derived exosomes and secreted DNA, we performed a proximity ligation assay (PLA) using these antibodies (**Fig. S7a**), in cryosections prepared from mouse ears obtained at 12 h post-TPA treatment, corresponding to the peak of neutrophil infiltration in skin. Incubation of cryosections with primary antibodies directed against either FLAP or dsDNA (but not both), showed a modest PLA signal in the form of random non-specific large spots throughout the TPA-treated (dorsal) side of PBS-injected mouse ear, likely a reflection of an association between the DNA probes and dead cell debris in the tissue (**Fig. S7b**). In sharp contrast, a robust PLA signal was observed in cryosections from PBS-injected mice when using both primary antibodies. As expected, the PLA signal is present in the dermis, is robust on the dorsal side of TPA-treated ear skin, and its intensity is dramatically decreased in the TPA-naïve ventral side of the dermis (**Fig. 7a, b**). Importantly, no specific PLA signal is observed in the dermis of TPA-treated mice that had been pre-treated with DNase I (**Fig. 7a, b**). Quantification of the intensity of the PLA signal, and the Ly6G and citrullinated histone H3 immunostaining patterns in 12 h TPA-treated skin sections of PBS-injected mice revealed a similar enrichment at sites proximal to the TPA treatment (dorsal side) (**Fig. 7c**) and is consistent with the co-secretion of LTB_4_ exosomes and DNA, proximal to the sites of inflammation. Collectively these findings establish that neutrophils infiltrating sites of inflammation release **S**ecreted **E**xosome-**A**ssociated **D**NA (SEADs) capable of modulating inflammatory responses.

**Figure 7.**
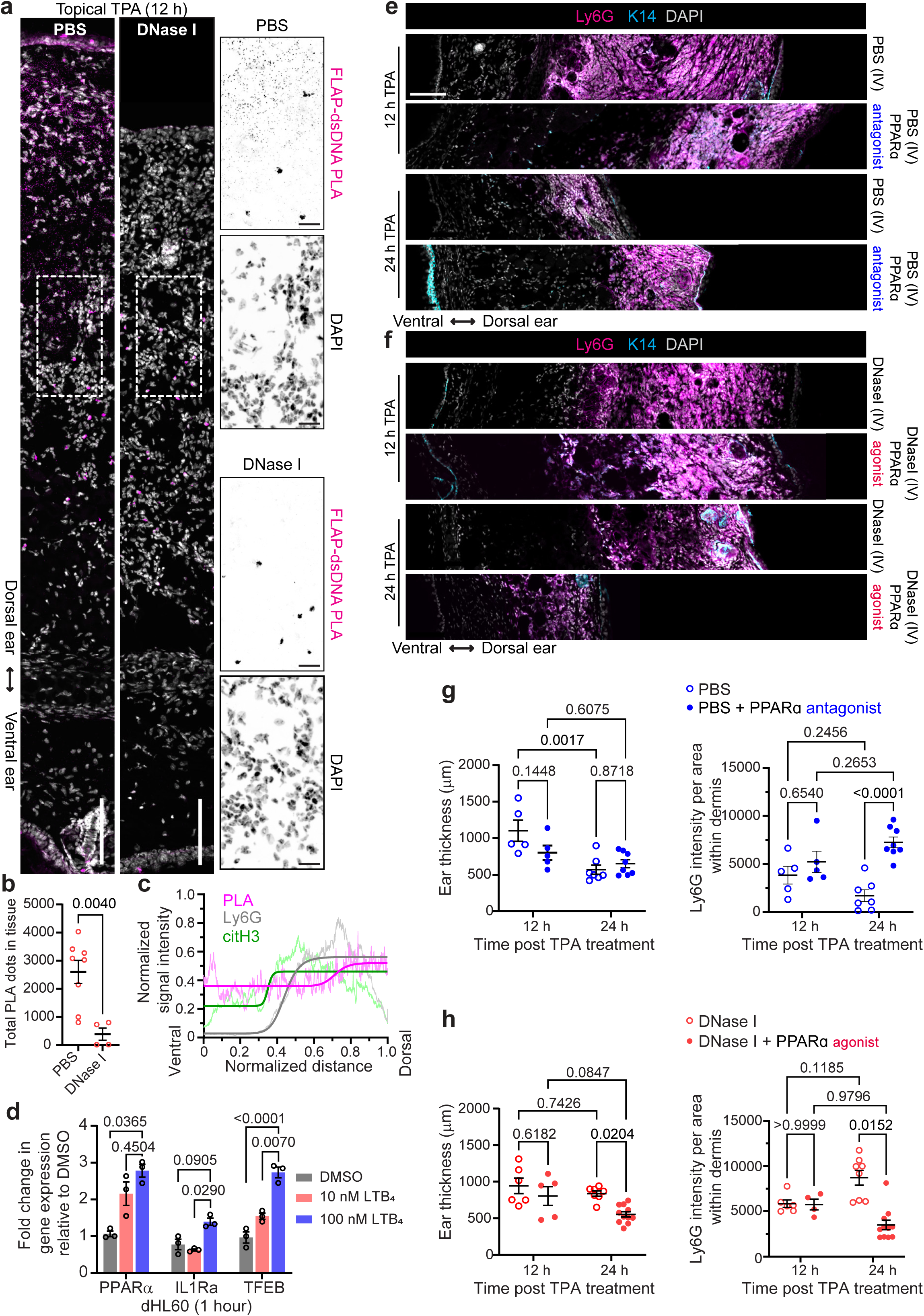
Disruption of SEADs impedes neutrophil resolution during TPA-induced ear inflammation in a PPARα-dependent fashion. **a.** Representative Airyscan microscopy images of TPA-treated (12h) ear cryosections (20 µm) from mice injected with either 100 μg DNase I or PBS, showing FLAP-dsDNA PLA signal (magenta dots) and DAPI (gray, nuclei). Images are presented as a ‘sum of slices’ projection of z-stacks. The scale is 100 μm and it is 20 µm in the insets. N=3. **b.** Scatter dot plot showing total FLAP-dsDNA PLA dots in the ear post 12 h TPA treatment. The maximum intensity projection images of 20 µm cryosections were calculated by object segmentation using CellProfiler and data points (red circles) pooled from 3 independent experiments are presented as mean ± s.e.m. P values calculated using the Mann-Whitney test are shown. **c.** Graph showing the four-parameter logistic curve fit (thick lines) of normalized PLA, Ly6G, and citH3 intensity distribution from the ventral to the dorsal side in inflamed ear post 12 h TPA treatment. The wavy lines represent the mean of 7 datapoints pooled from 3 independent experiments. **d.** Bar graph showing the fold change in the expression of various PPARα-responsive genes in dHL60 cells. Datapoints (black open circles) are plotted as mean ± s.e.m. and P values quantified using two-way ANOVA are shown. N=3. **e-f.** Representative Airyscan microscopy images of 20 µm thick cryosections of TPA-treated ear under different conditions as mentioned on the right side of the respective panels showing the relative localization of Ly6G (magenta, neutrophils), K14 (cyan, epidermis), and DAPI (gray, nuclei). The tiled z-stacks are presented here as a ‘sum of slices’ projection. The scale is 100 μm. N=3. **g-h.** Scatter dot plots showing the changes in ear thickness and Ly6G intensity per unit area within the dermis of inflamed ears under different conditions at 12 and 24 hrs post-TPA treatment, as shown in panels e-f. Datapoints (circle/dots) pooled from 3 independent experiments are plotted as mean ± s.e.m. and P values calculated using two-way ANOVA analysis are shown on the graph.

### The resolving effects of SEADs on acute inflammation require PPARα activation

It has been reported that high nanomolar concentrations of LTB_4_ promote BLT1 endocytosis^75^, and activate BLT2 and PPARα, both of which exert anti-inflammatory effects^18,26,27,76,77^. Indeed, as expected, we found an increase in the expression of PPARα-responsive genes, PPARα^78^, IL1Ra^79^, and TFEB^80^, in dHL60 cells upon treatment with LTB_4_. Notably, the transcriptional response was stronger for IL1Ra and TFEB at high (100 nm) compared to low (10 nM) LTB_4_ concentrations (**Fig. 7d**). From this we posit that (i) the clustering of LTB_4_-containing exosomes within SEADs leads to a local spike in LTB_4_ concentration that triggers the resolution of inflammation through BLT2/PPARα signaling, and that (ii) inefficient PPARα activation is responsible for the delay in this resolution, observed in the presence of DNase I. Accordingly, we investigated the role of PPARα activation as a contributor to the SEAD-mediated resolution of inflammation in TPA-treated mouse ear skin. We found that the topical application of GW6471^81^, a PPARα-specific antagonist^82^, to mouse dorsal ear skin at 6 h post-TPA treatment in PBS-injected mice results in a defective clearance of infiltrated neutrophils 24 h after TPA treatment. Treatment with GW6471 did not, however, have a measurable impact on ear thickness 24 h post-TPA treatment (**Fig. 7e, g**). In striking contrast, topical application of a PPARα agonist^81^, WY14643^83^, to mouse dorsal ear skin at 6 h post-TPA treatment rescues both the excessive neutrophil infiltration and ear thickness phenotypes observed 24 h post-TPA treatment in DNase I-injected mice (**Fig. 7f, h**). Importantly, we found that treatment with the PPARα agonist and antagonist did not affect either ear thickness or neutrophil infiltration at 12 h post-TPA treatment (**Fig. 7e-h**), suggesting a role for PPARα in the resolving but not in the initial phase of inflammation, consistent with previous reports^18,29^. Overall, these findings strongly suggest that SEADs facilitate the initiation of neutrophil clearance during the resolution phase of sterile inflammation by eliciting PPARα activation.

## DISCUSSION

The findings reported in this study reveal that migrating neutrophils co-secrete LTB_4_-containing exosomes and nuclear DNA as SEADs through a regulated, non-lytic mechanism initiated at the nuclear envelope. Our data indicate that, unlike PMA-induced necroptotic cell death and suicidal NET release^49,51,52,84^, SEAD release is a rapid and repetitive process involving nSMase-mediated NE-budding and NE-derived MVB biogenesis. We show that the packaging of DNA within NE-derived MVBs requires LBR and is facilitated by chromatin decondensation. Finally, we show that SEADs regulate chemotaxis *ex vivo* and promote the resolution of neutrophil infiltration *in vivo.* As DNase I treatment disrupts SEADs and not the process of secretion *per se*, we propose that NE-derived exosomes ensconced within SEADs give rise to a spatiotemporal spike in LTB_4_ concentration at the site of inflammation, allowing for the activation of the anti-inflammatory transcription factor PPARα and the initiation of resolution^18,28,30,85^. Specifically, our findings provide mechanistic insight into previous studies reporting the presence of cytoplasmic vesicles filled with chromatin-like structures in *S. aureus*-infected neutrophils, which were proposed to participate in non-lytic NET release ^11,12^. Overall, our study unveils the unexpected role of extracellular DNA - secreted by infiltrating neutrophils - in promoting the resolution of inflammation. While the role of PPARα in promoting the resolution of inflammation has been widely reported, our work provides a mechanism for the timely activation of PPARα during acute inflammation. Disruption of SEAD-mediated signaling may lead to a delay in the resolution of acute inflammation and potentially disease-promoting chronic inflammation.

### The unique neutrophil nuclear composition favors NE-budding and vital DNA secretion

The mechanisms mediating the presence of nuclear DNA within NE-derived MVBs reported here differ from the mechanisms believed to regulate nuclear content loading in ILVs within conventional CD63-positive MVBs generated in cancer cells^86^. We reason that this is partly due to the unique nature of neutrophils, which are non-dividing cells and thereby lack micronuclei^87^, a prerequisite for DNA packaging within conventional MVBs^86^. Moreover, the limiting membrane of NE-derived MVBs is decorated with LBR and lacks the core NE protein lamin A, which along with LAP2α and emerin is present in hybrid organelles formed upon fusion of micronuclei with CD63-positive MVBs during DNA loading in conventional exosomes^24,86,88^. Furthermore, in contrast to rupture-prone micronuclei enriched with core NE proteins^89^, the unique composition of the chromatin-containing NE-derived MVBs may instill structural integrity in these vesicles and the presence of ESCRT components^24^ may prevent spontaneous membrane rupture. Indeed, while we have not yet ruled out the possibility of cGAS-STING pathway activation^90,91^ in chemotaxing neutrophils, the orientation of the chromatin-bound Tudor domain of LBR towards the lumen of NE-MVBs and the disappearance of histone H3 signal in DNase I-treated exosomes strongly suggest that DNA is encapsulated within the lumen of NE-MVBs, and therefore protected from cytosolic cGAS. In addition, the absence of LBR in the exosomes provides evidence that the secreted nuclear DNA is most likely non-specifically associated with the exosomes. The absence of LBR – a cholesterol synthesizing enzyme^33^ – from the NE-derived exosomes is potentially due to the enrichment of ceramide on these exosomes and on lipid-ordered NE-membrane microdomains that are crucial for NE-derived exosome biogenesis^24^, and to the ability of ceramide to displace cholesterol-rich regions from these lipid-ordered microdomains^92^.

In human neutrophils, heterochromatin is uniquely organized along the NE and undergoes rapid decondensation following activation^61^, and migration through constricted spaces^93^. Not only does rapid chromatin remodeling facilitate nuclear softening and enhance neutrophil migration^94^, but it also initiates expedited inflammatory gene expression in response to neutrophil activation^95,96^. Consistent with the role of chromatin decondensation during ionophore (A23187)-induced NET release^62^, we found that LTB_4_ rapidly increases histone H3 acetylation, indicating chromatin decondensation, which we show is crucial for DNA secretion from chemotaxing neutrophils. The 5LO-positive NE-derived MVBs feature histone H3 acetylated at lysine K27, a modification site pivotal for euchromatin remodeling^96–98^. In addition, the fact that we find an increase in the percentage of DNA-secreting cells upon HDAC inhibition suggests that DNA decondensation is a limiting factor for DNA packaging and release. Though long-term and stable chromatin modifications such as histone H3 methylation are required for efficient heterochromatin remodeling during neutrophil differentiation^99^, the slow kinetics of histone methyltransferases and demethylases^100^ render it improbable that changes in histone methylation levels induce the rapid chromatin remodeling required during DNA secretion from chemotaxing neutrophils.

Observations across multiple healthy donors revealed that the percentage of DNA-secreting neutrophils saturates at ∼50 percent during one hour of migration. This can be attributed to either the dispersive loss of chemoattractant gradient^38,39^ or an unidentified auto-inhibitory regulatory mechanism. Alternatively, this may suggest the existence of a neutrophil subpopulation predisposed to DNA secretion. Interestingly, our finding that the nucleus of the DNA-secreting neutrophils is less lobulated than the nucleus of cells devoid of DNA secretion is consistent with a previous report showing elevated NET release from immature low-density neutrophils (LDNs) upon ionophore stimulation^101^. LDNs are also characterized by aberrant, ring-shaped nuclei with poor spatial segregation of heterochromatin and euchromatin^102,103^.

### ‘SEAD’ing is distinct from suicidal NETosis and cell death

We show that the repetitive secretion of DNA from chemotaxing neutrophils is a regulated event, that is not a consequence of cell death, but rather a result of nSMase-mediated NE budding. The observation that inhibition of ferroptosis by ferrostatin 1 tends to reduce DNA secretion could be related to the previous finding that ferrostatin 1 suppresses the activity of lipoxygenases^104^, the enzymes crucial for eicosanoid biogenesis. Given that cholesterol exerts a protective effect on ferroptosis^105^ and that LBR modulates cholesterol biosynthesis^33^, the high expression of LBR in neutrophils may be crucial for safeguarding activated neutrophils from ferroptotic cell death resulting from elevated lipoxygenase activity. Of note, while LTB_4_ signaling increases calcium flux in zebrafish neutrophils transiently and repetitively^106^, treatment with an ionophore (A23187) leads to a saturating and persistent increase in intracellular calcium that initially promotes LTB_4_-biogenesis^107^, and NET release^62^. However, the persistent A23187-mediated increase in calcium later activates the ferroptotic cell death pathway^108^. A similar persistent calcium increase dependent on PKC activity has been observed for PMA-induced suicidal NET release^48,49^ (“NETosis” – a terminology reserved for NET release associated with cell death^109^) and necroptotic cell death^84^. PMA-induced suicidal NETosis requires the cleavage of histones mediated by the nuclear translocation of myeloperoxidase and neutrophil elastase, which promote chromatin decondensation and nuclear swelling followed by cell lysis during DNA secretion^110^. We envision that neutrophils at sites of uncontrolled inflammation during severe infection, exhibit persistent calcium increase leading to necroptotic/ferroptotic/pyroptotic cell death^111^ and subsequent suicidal NETosis^112^. While NETosis aggravates inflammation^113,114^, our findings suggest that during acute sterile inflammation, infiltrating neutrophils release SEADs to promote the initiation of its resolution.

### SEADs facilitate LTB_4_ dual signaling during inflammation

We observed a peak in extracellular DNA levels in the dermis of inflamed skin 18 h after TPA treatment – a time point that coincides with the initiation of neutrophil clearance from the tissue. DNase I pre-treatment gave rise to a significant decrease in neutrophil clearance, monocyte infiltration, as well as co-occurrence of FLAP (NE-derived exosomes) and dsDNA (secreted DNA), in TPA-treated mice ear. These findings suggest a role for extracellular DNA-dependent LTB_4_ signaling in mediating the response of immune cells to acute inflammation. Indeed, the phenotypic manifestations we observed can readily be explained by the well-characterized pro- and anti-inflammatory effects of LTB_4_^115,116^. We envision that the transition is mediated through a switch from BLT1 to BLT2 signaling as sustained higher levels of LTB_4_ promote BLT1 endocytosis^75^. Interestingly, findings showing that high LTB_4_ concentration gives rise to a stepwise phosphorylation of BLT1 that promotes the activation of distinct signaling pathways suggest that signaling through BLT1 could also be involved in mediating anti-inflammatory effects^124^. High concentrations of LTB_4_ have been reported to induce neutrophil elastase release, thereby increasing endothelial permeability, which facilitates reverse migration of infiltrated neutrophils^85^. The anti-inflammatory transcription factor, PPARα has also been reported to play an integral role in lipid-mediator class switching from pro-inflammatory LTB_4_ to pro-resolving Lipoxin A4^117–120^, which is crucial for neutrophil clearance by reverse transmigration^121–123^. Our model, in which DNA secreted from infiltrating neutrophils captures the co-secreted LTB_4_-containing exosomes leading to a local spatiotemporal spike in LTB_4_ concentration, therefore provides a mechanism by which the anti- and pro-inflammatory effects of LTB_4_ are manifested during acute responses.

### Contrasting roles of secreted DNA in human pathology

The presence of extracellular DNA is an evolutionarily conserved phenomenon that has been observed across diverse life forms, including but not limited to humans^8,101,125,126^, cyanobacteria^127^, plants^128^, and social amoeba, *D. discoideum*^129^. While the excessive presence of secreted DNA/NETs has been mostly associated with adverse outcomes during various infections and hyper-inflammatory disorders^8^, recent studies suggest otherwise for certain conditions^13,14,16^. Aggregated NETs play a role in the spontaneous resolution of gout by breaking down inflammatory cytokines^14,16^. In contrast to the inflammatory nature of PAD4-dependent NETs^130^, NET release from PAD4-deficient mice promotes the upregulation of M2b macrophages and induces post-infarction resolution thereby increasing survival^13^. DNase therapy disrupts NETs in certain inflammatory conditions such as cystic fibrosis^131^, idiopathic bronchiectasis^132,133^, and sepsis^134,135^ but only yields short-term benefits and at times even adverse outcomes. Together, these observations necessitate a re-evaluation of the role of extracellular DNA secretion *in vivo* during sterile versus infectious inflammation, as a deeper understanding of the mechanisms controlling the resolution of inflammation and the regulatory functions of SEADs and/or extracellular DNA is crucial for the development of targeted approaches to treat inflammatory diseases associated with excessive DNA secretion.

## ACKNOWLEDGMENTS

We are thankful to the past and present members of the Parent and Coulombe laboratories for advice and support. We thank the Platelet Pharmacology and Physiology Core at the University of Michigan for providing human blood from healthy volunteers, the proteomics resource facility and Dr. Venkat Basrur for assistance with mass spectrometry data acquisition and analysis, and the microscopy core facility for assistance with TEM sample processing. We also thank P. Hanson (University of Michigan) for valuable suggestions. This work was supported by funding from the University of Michigan School of Medicine (C.A.P.), postdoctoral (916874) (S.B.A.), and predoctoral (AWD025905) (S.P.C.) fellowship awards from an American Heart Association, a Life Sciences Institute cubed award, the Arnold and Mabel Beckmann Foundation award to the University of Michigan Cryo-EM facility and by several grant awards from National Institute of Health, namely: T32 training program in Cell and Molecular biology GM145470 (S.P.C), T32 training program in translational research GM141840 (M.F.), GM150019 (S.M.), R01AR079418 (P.A.C.), and R01AI152517 (C.A.P.).

## AUTHOR CONTRIBUTIONS

C.A.P. and S.B.A. conceptualized the study and designed experiments with input from all authors. P.A.C. and S.M. conceptualized the *in vivo* and cryo-CLEM experiments, respectively. S.B.A. conducted most of the experimentation, S.P.C. conducted chemotaxis assays, Y.X. conducted the mouse experiments, and M.F. and S.P.C. conducted the cryo-CLEM experiments. S.B.A and S.P.C collected most of the data and performed the analyses, and J.Z.S. generated an initial pipeline for live imaging analysis, optimized further by S.B.A. M.F. and S.M. analyzed the cryo-CLEM data. S.B.A., C.A.P., and S.P.C. verified all experimental results and their analysis. S.B.A. and C.A.P. led the effort for writing the initial draft of the manuscript, with input from S.P.C. and P.A.C. S.B.A. prepared the figures with input from C.A.P., S.P.C., and P.A.C.

## METHODS

### KEY RESOURCES TABLE

| REAGENT or RESOURCE | SOURCE | IDENTIFIER |
| --- | --- | --- |
| **Antibodies** |  |  |
| Rabbit anti-5-lipoxygenase | Abcam | ab169755 |
| Mouse anti-5 lipoxygenase | BD Bioscience | 610695 |
| Rabbit anti-FLAP | Abcam | ab85227 |
| Mouse anti-histone H3 | Santa Cruz Biotechnology | 517576 |
| Mouse anti-histone H3 | Sigma-Aldrich | SAB4200651 |
| Rabbit anti- histone H3 (citrulline R2 + R8 + R17) | Abcam | ab5103 |
| Rabbit anti-histone H3 (K27 acetyl) | Cell Signaling Technology | 8173S |
| Rabbit anti-lamin B receptor | Abcam | ab32535 |
| Rabbit anti-lamin A/C | Proteintech | 10298-1-AP |
| Rabbit anti-lamin B1 | Cell Signaling Technology | 13435S |
| Rabbit anti-lamin B2 | Cell Signaling Technology | 12255S |
| Mouse anti-CD63 | BD Biosciences | 556019 |
| Mouse anti-GAPDH | Santa Cruz Biotechnology | 166574 |
| Mouse anti-dsDNA | Santa Cruz Biotechnology | 58749 |
| Chicken anti-keratin 14 | BioLegend | 906004 |
| Alexa647 conjugated rat anti-Ly6G | BioLegend | 127608 |
| Alexa561 conjugated rat anti-F4/80 | Invitrogen | 505-4801-80 |
| Alexa488 conjugated goat anti-rabbit IgG (H+L) | Invitrogen | A11008 |
| Alexa568 conjugated goat anti-mouse IgG (H+L) | Invitrogen | A11031 |
| **Experimental models: Cell lines and mouse models** |  |  |
| HL60 | ATCC | CCL-240 |
| HEK 293T | ATCC | CRL-11268 |
| FVB mouse (male) | Charles River Laboratories | 207 |

| **Chemicals, peptides, and recombinant proteins** |  |  |
| --- | --- | --- |
| Paraformaldehyde | Electron Microscopy Sciences | 15714 |
| Polybrene | Sigma-Aldrich | H9268 |
| SEA-Kem agarose | Lonza | 50004 |
| Acrolyl x-SE | Thermo Fisher Scientific | A20770 |
| Acrylamide | Sigma-Aldrich | A4058 |
| Bis-acrylamide | BioRad Laboratories | 1610156 |
| Sodium acrylate | Sigma-Aldrich | 408220 |
| VA-044 | Tokyo Chemical Industry | A3012 |
| Ammonium persulfate | Sigma-Aldrich | A3678 |
| Proteinase K | New England Biolabs | P8107S |
| Guanidine hydrochloride | Acros Organics | 50-01-1 |
| DNase I | ProSpec | ENZ-319 |
| SYTOXgreen | Thermo Fisher Scientific | S7020 |
| SYTOXorange | Thermo Fisher Scientific | S11368 |
| CellMask orange | Invitrogen | C10045 |
| MitotrackerRed CMXros | Invitrogen | M7512 |
| Fluo4-AM | Invitrogen | F14217 |
| Hoechst 33342 | Invitrogen | H21492 |
| Normal goat serum | Sigma-Aldrich | G9023 |
| Saponin | Sigma-Aldrich | 84510 |
| Phorbol myristate (PMA/TPA) | Cell signaling Technology | 4174 |
| Leukotriene B<sub>4</sub> | Cayman chemicals | 71160-24-2 |
| GSK484 | Cayman chemicals | 17488 |
| GSK2795039 | MedChem Express | HY-18950 |
| GW4869 | Sigma-Aldrich | 6823-69-4 |
| GSK-872 | MedChem Express | HY-101872 |
| Z-DEVD-FMK (caspase 3 inhibitor) | ApexBio | A1920 |
| Ferrostatin 1 | Cayman chemicals | 17729 |
| Optiprep | Sigma-Aldrich | D1556 |
| Anacardic acid | Fisher Scientific | 50-193-3872 |
| Trichostatin A | Calbiochem | 647926 |
| MK886 | Cayman chemicals | 10133 |
| **Critical commercial assays** |  |  |
| Duolink® In Situ PLA® Probe Anti-Rabbit PLUS, Affinity purified Donkey anti-Rabbit IgG (H+L) | Sigma-Aldrich | DUO92002 |
| Duolink® In Situ PLA® Probe Anti-Mouse MINUS, Affinity purified Donkey anti-Mouse IgG (H+L) | Sigma-Aldrich | DUO92004 |
| Duolink® In Situ Detection Reagents Red | Sigma-Aldrich | DUO92008 |
| CytoQuant LDH cytotoxicity assay | Invitrogen | C20301 |
| TMT10plex Isobaric label reagent and kits | Thermo Scientific | 90110 |
| **Software and algorithms** |  |  |
| Zen black | Zeiss | v2.3 |
| Fiji | ImageJ | v1.53q |
| Prism | GraphPad | v10.1.0 |
| Matlab | MathWorks | R2021b |
| CellProfiler | Broad Institute | v4.2.5 |
| Flowjo | Becton Dickinson & Co | v10.9.0 |
| Adobe Illustrator | Adobe | v26.3.1 |

### EXPERIMENTAL MODEL DETAILS

#### Cell lines

Human myeloid leukemia-derived promyelocytic cell line, HL60 (also known as PLB-985 cells) was obtained from ATCC (catalog #CCL-240) and cultured in IMDM medium (Gibco™ #12440053) containing 10% heat-inactivated fetal bovine serum (HI-FBS, Gemini Bioproducts #100-106), 20 mM HEPES buffer (pH 7.4), and penicillin-streptomycin antibiotic (Thermo Fisher Scientific #15-140-122) solution at a density of 0.2 to 1 million/ml. We differentiated HL60 into neutrophil-like cells by culturing them in IMDM containing 1.3 % DMSO, ITS (Insulin-Transferrin-Selenium) solution (Gibco™ #4140045) and 5% HI-FBS for 4 days. Media was replaced on day 2 as per previous protocols^136^. HEK293T cells, procured from ATCC (CRL-3216), were cultured in DMEM (Gibco™ #11965092) supplemented with 10% HI-FBS and used for lentivirus production to create stable HL60 cell lines.

Human neutrophils were isolated from the peripheral blood obtained by venipuncture of anonymous healthy donors recruited by the Platelet, Pharmacology, and Physiology Core facility at the University of Michigan. The blood collection procedure adhered to the institutional review board protocol (IRB #HUM00107120), approved explicitly for supplying de-identified blood for research. Consequently, we do not have any access to HIPAA information. All participants consented to provide their blood for research purposes and were financially compensated. All the de-identified healthy blood donors verified no intake of non-steroidal anti-inflammatory drugs for 48 hours before blood collection. Neutrophils were isolated using the previously described protocol^137^. In brief, whole blood was incubated with an equal volume of 3% dextran (Sigma-Aldrich #D1037) in 0.9% NaCl for 30 min at 37° C to facilitate the sedimentation and removal of red blood cells. The supernatant containing platelets, monocytes, lymphocytes, and neutrophils, was collected as plasma in a 50 ml centrifuge tube. The plasma was underlaid with one volume of Histopaque-1077 (Sigma-Aldrich #1077) and centrifuged at 400g for 20 min at room temperature to separate peripheral blood mononuclear cells from neutrophils obtained as cell pellets. Residual red blood cells were removed from neutrophils using ACK lysis buffer (Thermo Fisher #A1049201). This protocol yields >99% live neutrophils with close to 95% purity. Freshly isolated human PMN were resuspended in Hank’s Balanced Salt Solution (HBSS, Gibco™ #24020117) supplemented with 25 mM glucose (mHBSS) at a maximum cell density of 10^7^ cells/ml in a BSA-coated centrifugation tube.

#### Mice

All mouse experiments involved 12 to 16-week-old male mice in the FVB background and were approved by the Institutional Animal Use and Care Committee of the University of Michigan Medical School. Female mice were not used to avoid variable immune responses due to hormonal changes.

### METHOD DETAILS

#### Generation of CRISPR cas9 knockouts

To generate lentivirus, HEK293T cells were transfected with a mix of plasmids: VSV-G, psPAX2, and pLentiCRISPR V2 vector harboring SCR/LMNA/LBR single guide RNA (sgRNA). These were transfected at a 1:2:4 w/w ratio using Lipofectamine 3000 (Invitrogen #L3000001), according to the manufacturer’s instructions. Lentiviral particles harvested 48- and 72-hours post-transfection were pooled and concentrated using 4X virus precipitation solution (40% PEG 8000 and 1.2 M NaCl in PBS)^138^. Concentrated virus particles solubilized in IMDM were applied to HL60 cells in the presence of 8 μg/ml polybrene (Sigma-Aldrich #H9268). Clones expressing sgRNAs were selected with 2 μg/ml puromycin, with successful integration confirmed using western blotting and genetic sequencing. The LMNA sgRNA sequence: GCGCCGTCATGAGACCCGAC, and LBR sgRNA sequence: ATACCCATCCAATCAATCCG, were cloned into the pLentiCRISPR V2 plasmid, which was generously provided by the Zhang lab.

#### LDH release assay

We evaluated the integrity of the neutrophil cell membrane as a metric for cell death using CyQUANT™ lactate dehydrogenase (LDH) cytotoxicity assay (Invitrogen #C20300), according to the manufacturer’s instructions. Briefly, PMNs were plated on BSA-coated 12-well plates at a density of 1 x 10^6^ cells/ml in mHBSS. Cells were treated with either DMSO (vehicle), LTB_4_ (100 nM), or PMA (20 nM) for 2 h at 37 C. Cells treated with 1x cell lysis buffer were used as positive control and measure total (max.) cellular LDH. Post incubation, 50 µL culture supernatants were aliquoted into a 96-well plate and mixed with an equal volume of LDH reaction mixture. Following 30 min incubation at room temperature, shielded from direct light exposure, the color reaction was halted by adding the provided stop solution. Using SpectraMax M5 Multi-Mode Microplate Reader (Molecular Devices), the absorbance was quantified at both 490 nm and 680 nm (plate correction). The obtained data was corrected for background noise (plate) and nonspecific absorbance (mHBSS) and the level of LDH release was calculated and plotted as a percentage relative to maximal LDH release.

#### Nuclei isolation and histone extraction

To inhibit protease activity, the cells (10^8^/ml) were pretreated with 2 mM Pefabloc SC Plus (Roche #11873601001) for 15 min at 37 °C with rotation set to 10 RPM. Cells were then pre-treated with either DMSO as a control or HAT (200 nM Anachardic acid, Fischer Scientific #50-193-3872) or HDAC (400 nM Trichostatin A, Sigma-Aldrich #T1952) inhibitors for 30 min at 37 °C and stimulated with 100 nM LTB_4_ (Cayman chemicals #20110) for 5 min with rotation set to 10 RPM. The cells were centrifuged at 500g for 2 min at 4 °C, and the cell pellet was resuspended at a concentration of 50 x 10^6^ cells/ml in ice-cold hypotonic lysis buffer containing 10 mM HEPES pH 7.5, 4 mM MgCl_2_, 25 mM NaCl, 1 mM Dithiothreitol (DTT), and 0.1% NP-40, and lysed with 15 strokes of trituration^24^. Following centrifugation at 16,000g for 15 seconds at 4 °C, the nuclear pellet was resuspended in 0.2 N HCl in a volume required to maintain a density of 25 x 10^6^ nuclei/ml and incubated at 4 °C for 16 hours at a rotation set to 10 RPM^139^. The acid-extracted histones were collected in the supernatant fraction after centrifugation at 650g for 10 min at 4 °C and were neutralized with 2 M NaOH at 1/10 the volume of the supernatant before quantifying protein concentration using BCA^®^ Protein Assay kit (Thermo Scientific #23225)^139^. An equal amount of protein was boiled in 1x XT sample buffer at 95 °C for 10 min and electrophoresed on 4-12% Bis-Tris gel followed by electrotransfer onto PVDF membrane and was probed using antibody against: histone H3 K27Ac (1:500, Cell signaling technology #8173S), citrullinated histone H3 (1:2000, Abcam #5103), and histone H3 (1:1000, Sigma-Aldrich # SAB4200651).

#### Sub-cellular fractionation

dHL60 cells (100 million) were resuspended in mHBSS at a density of 10^7^ cells/ml and pretreated with 2 mM Pefabloc SC Plus (Roche #11873601001) for 15 min at 37 °C with rotation set to 10 RPM in a BSA-coated centrifugation tube. The cells treated with 100 nM LTB_4_ for 30 min at 37 °C on rotation set to 10 RPM were centrifuged at 400g for 2 min at 4 °C. The cell pellet was resuspended in disruption buffer (100 mM KCl, 3.5 mM CaCl_2_, 1.5 mm EGTA, and 10 mM PIPES pH 7.2) at a density of 10^7^ cells/ml. Controlled lysis of cells was performed by 3x freeze-thaw cycles in liquid nitrogen and room temperature water (10 min each). Disrupted cells were centrifuged at 400g for 20 min at 4 °C to pellet nuclei and intact cells. Supernatants were collected and centrifuged at 15,000g for 15 min at 4 °C to remove the mitochondrial pellet. The supernatant (1 ml) was collected and mixed with Halt™ protease inhibitor cocktail (ThermoFisher #78430) and overlayed on an 11 ml column of 5-30% linear gradient of iodixanol (Optiprep™, Sigma-Aldrich #D1556) in a 13 ml ultracentrifuge tube (Beckman Coulter #C14277). The linear gradient was generated by 3x freeze-thaw cycles of 20% Iodixanol in disruption buffer, alternating between -30 °C and room temperature for 30 min each^140^. The linearity of the gradient was verified by measuring the absorbance at 340 nm of 14 fractions (750 μl each), collected from the top, and diluted 10,000x in water. The sample on the gradient column was centrifuged at 100,000g for 16 hours at 4 °C, and proteins from collected fractions were precipitated using DOC-TCA-Acetone precipitation. Briefly, following 10 min room temperature incubation in 75 μl of 0.15% sodium deoxycholate (Thermo Scientific #89904), the protein was precipitated in final 30% trichloroacetic acid 6.1 N (TCA, MP Biomedicals #196057) in ice for 30 min. Protein pellets concentrated at 16,000g for 5 min at 4 °C, were washed with 500 μl ice-cold acetone to remove TCA at 16,000g for 5 min at 4 °C and resuspended in 100 μl 1x XT sample buffer (Bio-Rad #1610791), boiled at 95 °C for 10 min. The samples were analyzed on 4-12% criterion XT Bis-Tris polyacrylamide gel (Bio-Rad #3450124) electrophoresis followed by electrotransfer onto PVDF membrane and were probed using an antibody against CD63 (1:500, DSHB #H5C6), FLAP (1 µg/ml, Abcam #85227), LBR (1:1000, Abcam #32535), and Histone H3 (1:500, Sigma-Aldrich #SAB4200651).

#### Underagarose chemotaxis and live imaging

The chemotaxis assay was conducted according to the protocol established by Saunders et al.^141^ Briefly, 8-well chambered dishes with #1.5 glass bottom were coated with 1% BSA in DPBS for 1 hour at 37 °C to provide a hydrophobic surface for amoeboid migration of neutrophils. The agarose gel was prepared as 0.5 % agarose in a 1:1 ratio of DPBS: mHBSS poured into the dish and left to set for 45 min (15 min room temperature followed by 30 min at 4 °C). To visualize DNA secretion from migrating neutrophils, SYTOX™green/SYTOX™orange nucleic acid stain (Invitrogen™ #S7020) was added to the agarose before solidification. Subsequently, two 1 mm diameter wells were carved 2 mm from each other. One of the wells was filled with 100 nM LTB_4_ or fMLF diluted in mHBSS, successfully creating a chemical gradient detailed by Afonso and colleagues^22^ at approx. 50 pM/μm. A total of 50,000 cells in 5 µl mHBSS, pretreated with indicated inhibitors or DMSO vehicle control for 30 min at 37 °C were plated in the remaining well, and incubated at 37 °C for 1 h to allow migration. For immunofluorescence staining cells were fixed and processed as described below. For live imaging, cells were stained for 15 min at 37 °C before being plated in an agarose well with respective markers of subcellular structures: CellMask orange (plasma membrane, 1 µg/ml, Invitrogen™ C10045), Fluo4-AM (cellular calcium, 2 µM, Invitrogen #F14201), MitoTracker™ Red CMXros dye (intact mitochondria, Invitrogen™ M46752), and Hoechst 33342 (nuclear DNA, 1 µg/ml, Invitrogen #H3570). The images were acquired every 30-second interval for up to 3 hours using a Zeiss 880 confocal microscope fitted with 20x objective and controlled using Zen black software. For higher resolution, images were acquired using a 40x water-immersion objective, on Zeiss 880 fitted with an Airyscan detector and were presented after Airyscan processing on Zen black software. For assessing cell migration parameters images of migrating cells were acquired using a Zeiss Colibri microscope fitted with a 10x objective controlled using Metamorph software.

#### Exosome isolation and treatment

Exosomes were isolated from PMNs or dHL60 cells using the protocol outlined by Thery et al^142^. At least 50 million PMNs and 100 million *LMNA* or *LMNA/LBR* KO dHL60 cells were stimulated with 100 nM LTB_4_ at 37 °C for 30 min in BSA-coated centrifuge tubes at 10^7^/ml either in the presence or absence of 1 µg/ml DNase I (Fisher Scientific #NC9007308). Cells were pelleted at 500 g for 5 min at 4 °C and supernatants collected in BSA-coated tubes were centrifuged again at 4,000g for 20 min at 4 °C to remove apoptotic cells and larger microvesicles. Collected supernatants were incubated with final 8% PEG-6000 (Bio Basic #PB0432) in 20 mM HEPES and 500 mM NaCl solution (pH 7.4), at 4 °C for 36 h followed by centrifugation at 4,000g for 1.5 hr^143^. The concentrated exosome pellet was resuspended in 1 ml of exosome resuspension buffer (250 mM sucrose in 20 mM Tris-Cl buffer pH 7.4) and was overlayed onto an optiprep™ discontinuous gradient and centrifuged at 100,000g for 16 hours at 4 C. The optiprep gradient was generated by layering 3 ml of 40% optiprep, onto 3 ml of 20% optiprep followed by 3 ml of 10% optiprep and 2 ml of 5% optiprep diluted in exosome resuspension buffer. The purified exosomes were collected as fractions 4-9 (Iodixanol density 1.083-1.142 g/ml) as described in Majumdar et. al.^25^, the volume was readjusted to 12 ml in 13 ml SW41Ti rotor-compatible ultracentrifuge tubes and spun at 100,000g at 4 °C for 1 h. The purified exosome pellet was either resuspended in mHBSS^++^ for DNase treatment or in 1x XT sample buffer (reducing) for western blot analysis and vortexed intermittently for 15 min. For DNase treatment, one-half of the resuspended exosomes were treated with 20 units/ml of DNase I and the other half with an equal volume of PBS (vehicle) at 37 C for 30 min, diluted with mHBSS^++^, and were centrifuged at 100,000g for 1 h at 4 C. Treated exosome pellet was lysed in 1x XT sample buffer and boiled at 95 °C for 10 min. The samples were electrophoresed on 4-12% Bis-Tris gel, transferred onto PVDF membrane with 0.2 µm pore size, and were probed using anti-histone H3 (1:500, Sigma-Aldrich #SAB4200651), anti-LBR (1:1000, Abcam #32535), anti-5LO (1:1000, Abcam #169755) and anti-FLAP (1 µg/ml, Abcam #85227) antibodies and were detected using chemiluminescence on photographic films.

#### Expansion microscopy

To ensure the structural preservation of polymorphonuclear neutrophils (PMNs) following expansion, we slightly modified the previously described protocols^144,145^. PMNs were allowed to migrate under a 3 ml agarose layer placed over a 22 × 22 mm glass coverslip (#1.5, BSA coated) within a 35-mm petri dish for 1 h at 37 °C, then fixed with 1 ml of paraformaldehyde (4% PFA) and 0.05% glutaraldehyde in PHEM buffer (60 mM PIPES, 25 mM HEPES, 10 mM EGTA, and 2 mM MgCl2, with the pH adjusted to 6.9) at 37 °C for 20 min. The agarose was gently removed post-fixation, and the cells underwent immunostaining with a saponin-based buffer as previously detailed. Following immunostaining, cells on the coverslip were rinsed three times with 1× PBS and subjected to crosslinking with 3 mM acryloyl-X, SE (Sigma-Aldrich #A20770) in 1× PBS at room temperature overnight. The subsequent removal of the excess crosslinker was performed with three 15-min 1× PBS washes at room temperature. For gelation, we placed an 80 μl droplet of the gelation solution—containing 8% sodium acrylate, 10% acrylamide, 0.1% bis-acrylamide, and 1% VA-044 initiator—onto the coverslip and kept it at 4 °C for 10 min. We then assembled the coverslip in a gelation chamber as per the method described by Truckenbrodt et al.^144^, adding 200 μl of the gelation solution before placing the setup within a humidified chamber at 37 °C for at least 2 hours to allow polymerization. After the gelation chamber was disassembled, we briefly rinsed the polymerized gel on the coverslip with 1× PBS to wash away any remaining unpolymerized solution. Digestion of the gel was carried out using a buffer solution of 50 mM Tris-Cl, 800 mM guanidine hydrochloride, 1 mM EDTA, and 0.5% Triton X-100, adjusted to pH 8.0. We added proteinase K (8 U/ml, NEB #P8107S) immediately before use and incubated the gel at 37 °C for 1 hour in a 35 mm dish. Post-digestion, we washed the gel three times in 2 ml of 1× PBS, each wash lasting 15 min, at room temperature. For gel expansion, we carefully transferred the gel into a larger 100 mm dish, incrementally submerging it in dilutions of PBS (1× to 0.5× to 0.02× to 0.01× PBS, 20 min each) at room temperature, followed by an overnight soak in deionized water (ddH2O) at 4 °C. The expansion multiplied the size of the gel approximately fourfold as measured with a ruler. We then cut the gel to fit into a 12 mm glass-bottom dish (35 mm) coated with poly-L-lysine (Sigma-Aldrich #P8920). To prevent drifting during imaging, we secured the gel to the dish with a gentle press from a soft brush and filled the surrounding space with 100 μl ddH_2_O to avoid polymer shrinkage. Imaging was conducted using a Zeiss 880 confocal microscope equipped with an Airyscan detector and a 63× (1.4 NA) oil objective. Airyscan resolution is approximately 120 nm laterally and 350 nm axially; thus, after 4× sample expansion, the enhanced effective resolution fell between 30-40 nm laterally and nearly 100 nm axially. The collected z-stack images were spaced 160 nm apart, which, after adjusting for expansion, equated to an effective 50 nm interval. All size measurements mentioned are accounted for after the 4× calibration. We clarified zoomed images of buds or vesicles through deconvolution, using a rapid iterative algorithm, which served both as the basis for presentation and data quantification.

#### Transmission electron microscopy

Neutrophils were stimulated with 100 nM LTB_4_ in mHBSS for 15 mins at 37 °C in 35 mm glass bottom dishes precoated with 25 μg/ml fibrinogen in DPBS. Cells were fixed in 2.5% glutaraldehyde in 0.1 M sodium cacodylate buffer pH 7.4 (CB, EMS #15960) overnight at 4 °C and washed three times at room temperature with 0.1 M CB for 5 mins each, followed by post-fixation using 1% Osmium tetroxide and 1% Potassium ferrocyanide in 0.1 M CB for 15 mins at 4 °C. Samples were washed thrice with 0.1 M CB at room temperature for 3 min each, followed by 3 washes with 0.1 M sodium acetate buffer (pH 5.2) for 2 min at room temperature. Fixed cells were stained *en bloc* using 2% Uranyl Acetate in 0.1 M Sodium Acetate (pH 5.2) overnight at 4 °C. After washing thrice in deionized water at room temperature, cells were stained with 20 mM of lead nitrate in 30 µM L-aspartic acid solution (pH 5.5) for 5 mins at 37 °C to enhance TEM image contrast of membrane or fibrous structures. The stained sample was washed in deionized water 4 times at room temperature followed by dehydration in increasing concentrations of ice-cold ethanol: 10%, 30%, 50%, 70%, 80%, 90%, and 95%, for 5 mins each at 4 °C and then twice with 100% ethanol for 5 mins at room temperature. Serial infiltration of Spurr’s Resin was performed in 100% ethanol at a 2:1 v/v ratio for 30 mins, 1:1 ratio for 1 hour, 1:2 for 4 hours, and finally in pure Spurr’s Resin overnight (17 hours), all at room temperature. Samples were embedded in slice capsules for 15 mins at room temperature, followed by polymerization for 6 hours at 70 °C, and then filled with absolute Araldite-Embed resin into the capsule for 18 hours at 70 °C. Thin 70 nm sections obtained using ultramicrotome were sputtered with 7 nm carbon nanoparticles using the Leica EM ACE600 instrument to enhance image contrast. Images were acquired on JEOL JEM-1400+ LaB6 Transmission Electron Microscope at 60 kV.

### Cryo-correlative light and electron microscopy (cryo-CLEM)

#### Sample preparation for cryo-CLEM

Neutrophils were stimulated with 100 nM LTB_4_ in mHBSS for 15 mins at 37 °C in 35 mm glass bottom dishes containing glow-discharged Au SiO_2_ R1/4 200 mesh grids (Quantifoil GmbH) precoated with 25 μg/ml fibrinogen in DPBS. SYTOX™green nucleic acid stain (Invitrogen™ #S7020) was added at a 500 nM concentration for the duration of LTB_4_ treatment. Following stimulation, the grids were back-side blotted and plunged frozen into liquid ethane at -185 °C using a Leica EM GP2 automatic grid plunger with the following specifications: blot time - 10s, Temperature - 37 °C and humidity - 90%. The plunge-frozen grids were clipped into autogrids and were stored in liquid nitrogen till further use.

#### Cryo-fluorescence microscopy

The quality of cells and grids was assessed in widefield mode on a Stellaris-5 cryo-confocal laser scanning microscope equipped with a cryo-stage and x50/0.9 numerical aperture objective (Leica Microsystems). Z-stacks of desired regions were acquired in confocal-scan mode with a 547 nm laser line (560-580 nm emission), 0.5 µm spacing spanning ± 7 µm of the focus plane, and an x/y pixel size of 110 nm. Additionally, a reflection image was also acquired at each Z plane to position the SYTOX™orange signal relative to the holey-carbon film for targeted tomography acquisition. Maximum intensity projections of Z-stacks were generated using the Leica built-in processing tools package (LAS X 4.5.0.25531). Subsequent correlations between confocal and TEM images were performed on the cross-platform correlative software Maps 3.20 (ThermoFisher Scientific) using the holey-carbon film as a guide for tomography tilt-series positioning.

#### Tilt-series data acquisition, processing, and segmentation

Plunge-frozen grids containing neutrophils were imaged on a 300kV Titan Krios G4i transmission electron microscope (ThermoFisher Scientific) equipped with a K3 direct electron detector and an imaging filter (Gatan Inc.) operated in counting mode. Dose-symmetric tilt-series^146^ were collected from –60° to +60°, starting at 0° at a 3° interval, at correlated areas using SerialEM software^147^. The magnification was set to a pixel size of 3.315 Å/pixel, and the total dose per tilt-series was 120e^−^/Å^2^. Data were collected with a slit width of 20eV. Tilt-series were aligned by patch tracking and reconstructed by weighted back projection implemented in IMOD^148^. 15 iterations of a ‘SIRT-like’ filter were applied. Reconstructed tomograms were imported into EMAN2^149^ and features of interest were segmented using the convolutional neural network-based segmentation implementation within. The segmentation was manually inspected and used for making movies in AMIRA (ThermoFisher Scientific).

#### Flow cytometry

dHL60 cells at a density of 1 x 10^6^ cells/ml were treated with either DMSO (control) or 100 nM LTB_4_ for 30 min at 37 °C in a BSA-coated centrifuge tube with rotation set to 10 RPM. After centrifugation at 400g for 2 min at 4 °C, cell pellets resuspended in 100 μl FACS buffer (phosphate buffer saline containing 1% BSA and 0.1% sodium azide) were treated with human TruStain FcX™ (Biolegend #422302) for 15 min on ice followed by incubation with rabbit anti-human LBR antibody (Abcam #32535, 1:100) for 30 min on ice. Following 2x washes with 500 μl FACS solution at 400g for 2 min at 4 °C, cells were stained using goat anti-rabbit IgG - AlexaFlour 488 (ThermoFisher #A11008, 1:500) in 100 μl FACS buffer, followed by 2x washes with FACS buffer, and fixation using 4% paraformaldehyde in PBS at room temperature for 15 min. Fixed cells were washed and resuspended in PBS before acquisition on BD Fortessa flow cytometer.

#### Quantitative PCR analysis

A minimum of 10 million dHL60 cells were treated with either DMSO, 10 nM LTB_4_, or 100 nM LTB_4_ in mHBSS for 1 h. Cells were lysed in Trizol™ reagent and total RNA was purified using RNeasy Mini kit (Qiagen # 74104). cDNA was generated from isolated RNA at a concentration of 1 μg per 10 μl reaction volume, using oligo(dT)_20_ primers provided within SuperScript™ IV First-Strand Synthesis System (ThermoFisher Scientific # 18091050) as per manufacturer’s instructions. The qPCR reactions of 10 μl each (in triplicates) using cDNA (100 ng per reaction) were amplified using PowerUp™ SYBR™Green Master mix (applied biosystems # A25742) and gene-specific primers (300 nM): PPARα (Forward-CTATCATTTGCTGTGGAGATCG, Reverse-AAGATATCGTCCGGGTGGTT), sIL1Ra (Forward-GCCTCCGCAGTCACCTAAT, Reverse-TCCCAGATTCTGAAGGCTTG), TFEB (Forward-CCAGAAGCGAGAGCTCACAGAT, Reverse-TGTGATTGTCTTTCTTCTGCCG), and 18S rRNA (Forward-GCTTAATTTGACTCAACACGGGA, Reverse-AGCTATCAATCTGTCAATCCTGTC. QuantStudio 5 system was used to monitor the cDNA amplification and determine cycle threshold (Ct) values. The relative change in gene expression was calculated using 2^-ddCt, with Ct values of β-actin as housekeeping gene control. Data was exported from Microsoft Excel and plotted in GraphPad Prism.

#### Mass spectrometry analysis

dHL60 cells were lysed in RIPA buffer (G-Biosciences # 786-489) on ice by sonication with 3 sec ON/OFF cycles for 10 cycles. An equal amount of protein (75 ug) quantified using BCA assay was processed for tandem-mass tagging (TMT) (FisherScientific™ #90110) according to the manufacturer’s instructions as described below.

#### Protein digestion and TMT labeling

Samples (75 ug per condition) were reduced in 5 mM dithiothreitol (DTT) for 30 min at 45 °C followed by alkylation with 2-chloroacetamide (15 mM) for 30 min at room temperature. The alkylated samples were precipitated at -20 °C overnight in 6 volumes of ice-cold acetone, followed by centrifugation to collect precipitated protein which was later air-dried and resuspended in triethylammonium bicarbonate (TEAB, 0.1 M) buffer. The resuspended proteins were digested in trypsin/Lys-C mix (1:25 protease: protein ratio; Promega) for 16 hours at 37 °C in a thermomixer. The TMT 10-plex reagents dissolved in 41 μl of anhydrous acetonitrile, followed by mixing entire digested protein samples and labeling the peptide at room temperature for an hour. The labeling reactions were quenched by incubating for 15 min in 8 μl of 5% hydroxyl amine. Labeled samples were pooled and vacuum dried, followed by offline fractionation of pooled samples (∼675 μg) that were resuspended in 675 μl of 0.1% trifluoro acetic acid (TFA). Two 100 μl aliquots were subjected to basic reverse phase spin column fractionation using two columns (Thermo Scientific # 84868) with elution pooled after fractionation. Fractions were dried and reconstituted in 9 μl of 0.1% formic acid and 2% acetonitrile mix for LC-MS/MS analysis.

#### Liquid chromatography-mass spectrometry analysis

To obtain superior quantitation accuracy, we employed multinotch-MS3 (McAlister GC)^150^ which minimizes the reporter ion ratio distortion resulting from fragmentation of co-isolated peptides during MS analysis. Orbitrap Fusion (Thermo Fisher Scientific) and RSLC Ultimate 3000 nano-UPLC (Dionex) were used for data acquisition. Briefly, 2 μl sample was resolved on a PepMap RSLC C18 column (75 μm; Thermo Scientific) at the flow rate of 300 nl/min using 0.1% formic acid/acetonitrile gradient system (2-22% acetonitrile for 150 min, 22-32% acetonitrile for 40 min, and 20 min wash at 90% acetonitrile followed by re-equilibration for 50 min) and directly sprayed onto the mass spectrometer using EasySpray source (Thermo Fisher Scientific). The mass spectrometer was optimized to collect one MS1 scan (Orbitrap; 120K resolution; AGC target 2x10^5^; max IT 100 ms) followed by data-dependent, “Top Speed” (3 seconds) MS2 scans (collision-induced dissociation; ion trap; NCE 35; AGC 5x10^3^; max IT 100 ms. For multinotch-MS3, the top 10 precursors from each MS2 were fragmented by HCD followed by Orbitrap analysis (NCE 55; 60K resolution; AGC 5x10^4^; max IT 120 ms, 100-500 m/z scan range).

#### Mass spectrometry data analysis

Using Proteome Discoverer (V2.4; Thermo Fischer) MS2 spectra were searched against the SwissProt human protein database (UniProt-filterd_Hsapiens_Reviewed_12132021.fasta) using the following search parameters: MS1 and MS2 tolerance set to 20 ppm and 0.5 Daltons, respectively; carbamidomethylation of cysteines (57.02146 Daltons) and TMT labelling of lysine and N-termini of peptides (229.162 Daltons) were considered for static modifications. While oxidation of methionine (15.9949 Da) and deamidation of asparagine and glutamine (0.98401 Da) were considered for variable modifications. Identified proteins and peptides were filtered to retain only those that passed ≤1% FDR threshold. The proteins were quantified using high-quality MS3 spectra with the following parameters: average signal-to-noise ratio of 10 and <50% isolation interference.

#### Mouse sterile inflammation model and drug treatments

Mice were treated with 20 μl of 0.2 mg/ml of TPA (Cell Signaling Technology #4174S), topically on the dorsal side of ear skin, with the other ear treated with an equal volume of acetone as vehicle control^67^. To disintegrate secreted DNA in the inflamed tissue, mice were injected intravenously in the tail with either 100 µl of PBS or recombinant human DNase I (PROSPECT #ENZ-319; 1 mg/ml) 1 h before TPA treatment. To perturb the PPARα pathway, the dorsal skin of the ear was treated topically with either 100 µg of PPARα agonist (WY14643, Sigma-Aldrich #C7081) or antagonist (GW6471, Sigma-Aldrich #G5045) dissolved in DMSO, 6 hours after TPA treatment for the remainder of the experiment^81^.

#### Tissue collection and cryosectioning

Treated mice were humanely euthanized by CO_2_, and their ear tissue was harvested at the designated time points. The samples were then embedded in the optical cutting temperature (O.C.T.) compound at -40 °C (Sakura® Finetek USA, #4583). With the CRYOSTAR NX50 Cryostat™ (ThermoScientific) set to -20 °C and using an MX35 ultramicrotome blade (Epredia #3053835), we prepared either 20 or 5 µm thick cross sections of the ear tissues. These sections were then transferred onto positively charged microscopic slides (VWR #48311-703). Until required for subsequent analysis, the slides bearing tissue sections were stored at -40 °C.

### Immunofluorescence staining, proximity ligation assay, and microscopy

#### For cells

Neutrophils migrating under the agarose for 1 h in 8 well chamber dishes were fixed using 4% paraformaldehyde (PFA, EMS #) in PHEM buffer for 20 min at 37 °C followed by agarose removal from the wells. Cells were washed thrice with PBS to remove residual PFA and permeabilized, blocked, and stained with primary antibodies against LBR (Abcam #32535, 1:400), histone H3 (Santacruz Biotech #517576, 1:100), 5LO (BD biosciences #610094, 1:200), and histone H3 K27Ac (cell Signaling Technology #8173T, 1:200), in staining solution (0.4% saponin, 2% goat serum in PHEM buffer) overnight at 4 °C. Cells were washed thrice with a staining solution for 5 mins each at room temperature, and incubated with secondary antibodies (Invitrogen, 1:500) against primary antibodies in staining solution for 30 min at room temperature. Cells were counterstained with Hoechst and Phalloidin to visualize nuclei and actin during secondary antibody incubation. Cells were washed thrice with PBS for 5 min each, and Immu-mount™ (Fisher Scientific #9990402) was added to cells before imaging under 63x oil-immersion objective in a Zeiss LSM 880 confocal microscope fitted with Airyscan detector. The images were acquired as z-stacks at an interval of 160 nm and were reconstructed using Airyscan processing in Zen 3.4 Black edition software.

#### For tissue

The frozen tissue section on glass slides was thawed and air dried for 10 min followed by fixation using 4% PFA in PBS for 10 min at room temperature. The tissue sections were washed thrice with PBS for 5 min each. A circle was drawn on the slide around the tissue with a hydrophobic barrier pen to generate a staining area, followed by blocking the tissue section for 30 min at room temperature with blocking buffer (2% normal goat serum and 1% BSA in PBS). The blocked tissue sections were stained using primary antibodies: chicken anti-keratin 14 (Biolegend #906004, 1:1000), rabbit anti-citrullinated histone H3 (Abcam #5103, 1:800), rabbit anti-FLAP (Invitrogen #MA5-37933, 1:200) and mouse anti-dsDNA (Santacruz Biotechnology #58749, 1:100) in blocking solution overnight at 4 °C. Following washes with PBS thrice for 5 min each, the cells were stained with secondary antibodies: goat anti-chicken Alexa568 (Invitrogen #A11041, 1:1000), goat anti-chicken Alexa647 (Abcam #15071, 1:1000), goat anti-rabbit Alexa488 (Abcam #150077, 1:1000), and goat anti-mouse Alexa647 (Invitrogen #A21241, 1:1000) in blocking solution for 1 h at room temperature. Neutrophils and macrophages were stained using Alexa647 conjugated rat anti-Ly6G (Biolegend #127609, 1:100) and Alexa561 conjugated rat anti-F4/80 (Invitrogen #505-4801-80, 1:100), respectively with the secondary antibody. The unbound secondary antibody was washed thrice with PBS for 5 min each and nuclei were stained with DAPI (1ug/ml) for 5 min in PBS followed by washing thrice with PBS. The samples were mounted using FluorSave™ Reagent (Millipore #345789), secured using a #1.5 coverslip, and air-dried overnight before image acquisition.

#### For proximity ligation assay

PFA-fixed tissue sections were incubated with anti-FLAP and anti-dsDNA antibodies overnight at 4 °C in antibody diluent followed by processing according to manufacturer’s instructions for Duolink^®^ proximity ligation assay (Sigma #DUO92101). Briefly, antibody-stained tissue was washed and incubated with PLA probes for 1 h at 37 °C followed by a ligation mixture for 30 min at 37 °C and then with an amplification mixture at 37 °C for 100 min. Tissue was then stained with DAPI (1 µg/ml) for 10 min at room temperature, washed, and mounted using FluorSave reagent. Imaging was performed on a Zeiss 880 microscope fitted with an Airyscan detector using a 40x water immersion objective. To image the full distal-to-ventral length of ear sections, tiled images with 512x512 resolution were acquired. To cover the full thickness of tissue sections z-stack at an interval of 600 nm was acquired.

#### Quantification and statistical analysis

Intensity profiles of RGB channels (Figs. 1b, d) across the diameter of the vesicle of interest were determined using the RGB profiler plugin (https://imagej.net/plugins/rgb-profiler) from the Fiji image analysis tool. The resulting data were exported to GraphPad prism and plotted as histograms with mean ± s.e.m. To quantify the spatial correlation between 5LO, LBR, and Hoechst in NE-derived buds/MVBs, the intensity of 3D images was thresholded using maximum entropy parameters, and Pearson’s R-value (Fig. 1f) was determined using the JaCoP plugin in Fiji image analysis software. For quantification of morphological features in DNA-secreting cells, CellProfiler-based cell segmentation analysis was used.

For parameter tracking of DNA-secreting neutrophils (Fig. 3, S2, S3), nuclei were segmented using the Otsu threshold option in the “IdentifyPrimaryObjects” module, which was used to identify cells based on the CellMask image using Watershed-Image option and otsu threshold in “IdentifySecondaryObjects” module in CellProfiler. Cells were shrunk 2 pixels and expanded 3 pixels, followed by subtracting shrunk objects with expanded objects, to obtain a 1.5 µm thick ring 1 µm outside plasma membrane. Size, shape, intensity, and intensity distribution of Hoechst (nuclear DNA), Fluo4-AM (intracellular calcium), and CellMask (for PM rupture) within the nuclei, cell, and cytoplasm were measured (Fig. 3b, S2 c-d, S3 a-e). The identified extracellular ring was assigned to respective cells using the “RelateObjects” module, and the presence or absence of SYTOXgreen signal in the extracellular ring was used to classify respective cells as positive or negative for DNA secretion. The percentage of DNA secretion-positive cells or of total cells that migrated during the observation window were plotted as shown in Fig. 3i, S2 f-g, 4e, and 5f. The SYTOXgreen signal was segmented using the Otsu threshold and was assigned as secreted DNA. The cytoplasm positive for SYTOXgreen signal was assigned as undergoing cell lysis as plotted in fig. S2d. The outlines of the segmented objects (Cell, nuclei, secreted DNA) were overlayed using the “OverlayOutlines” module and exported as an image, as shown in fig. 3a and S2b. All the cells were tracked throughout migration in multiple frames using the nearest neighbor algorithm with an average cell diameter of 10 µm and maximum intercellular distance to consider matches set to 60% of the max. length of migrating cells. The average speed before, after, and during DNA secretion was quantified by dividing the total distance traveled (calculated using the formulae below) by the time required to migrate through that distance (3f). Average directionality is calculated as the ratio of displacement to Euclidean distance (3g). A two-tailed correlation matrix between speed and directionality for indicated conditions was generated using GraphPad Prism software and is plotted in fig. 3h.

The nuclear form factor (fig. S4d) determined as the area to perimeter squared was quantified by nuclear segmentation using minimum cross entropy algorithm in the “IdentifyPrimaryObjects” module followed by the “MeasureObjectSizeShape” module of CellProfiler image analysis software. To determine Heterochromatin spots (fig. S4f) based on Hoechst 33342 intensity distribution, the spot features were enhanced using the “EnhanceOrSupressFeatures” module followed by masking with segmented nuclei. These masked objects were further segmented and identified as speckles using a minimum cross-entropy algorithm in the “IdentifyPrimaryObject” module. These speckles (heterochromatin spots) were related to the original nuclei and followed by shape and number analysis using the “MeasureObjectSizeShape” module. To calculate the LBR NE-cytoplasmic ratio (Fig. S4c), nuclei were expanded and shrunk ∼0.2 µm, and the shrunk nuclei were subtracted from expanded nuclei to get ∼400 nm thick nuclear envelope objects using “IdentifyTertiaryObjects” module. The ratio of LBR intensity in NE objects to the cytoplasm was quantified using Microsoft Excel and plotted using GraphPad Prism software. To identify nuclear invaginations (Fig. S4e), the LBR images were segmented using a minimal cross-entropy algorithm with strict parameters in the “IdentifyPrimaryObjects” module, to segment only the most intense signal. Previously identified NE objects were subtracted from segmented LBR objects using the “IdentifyTertiaryObjects” module to obtain LBR objects within the nuclei excluding NE. These objects were related to the corresponding cells and object (invaginations) number and were quantified using the “MeasuerObjectSizeShape” module.

Histone H3 intensity in the LBR-positive NE buds and vesicles (NE blebs) (Fig. 5e-f) was quantified using the “MeasureObjectIntensity” module. To segment NE buds and vesicles, the first the nuclei were segmented by applying minimum cross-entropy threshold on Hoechst 33342 images, using the” IdentifyPrimaryObjects” module, followed by segmentation of LBR positive objects by the otsu threshold of LBR images. Identified nuclei were expanded to 1 μm and were subtracted with LBR objects using “IdentifyTertiaryObjects” yielding LBR-positive blebs. Since LBR staining showed some non-nuclear/vesicular, possibly ER structures, the LBR-positive blebs were filtered based on diameter and area extent (>0.4) and identified as NE-blebs. The obtained H3 intensity was normalized to the area of NE blebs and plotted using a GraphPad prism. Detailed cell profiler pipelines with all the parameters used for object segmentation are published on the CellProfiler server.

Using Image J (Fiji) image analysis software, z-stack images of mice ear sections were converted to a “sum of slices” projection to preserve the intensity distribution along both xy- and z-axis, which are presented in Fig. 6e, S6, 7b, 7e, 7e, and S7. The thickness of the ear, and Ly6G and citH3 intensity across mice ear dermis were quantified by manually creating a region of interest (ROI) and applying the analyze>measure feature in Image J. Integrated intensity and area were exported and compiled in an Excel sheet and plotted using GraphPad prism. The intensity profile of PLA signal across the length of ear tissue from the sum of slices projection of z-stacks was quantified by first manually creating a rectangular ROI encompassing the entire tissue area and then using the analyze>plot profile feature of image J. The integrated intensity changes across the ear section were exported and analyzed in GraphPad Prism. The data was normalized in GraphPad prism within a scale of 0 to 1 to account for variation in PLA intensity among experimental replicates and different tissue sections, along with the variation in the thickness of ear tissue.

Analysis of cell migration features as presented in figures S5c-f and 6a-d was performed using the Trackmate plugin in the ImageJ platform on the Hoechst 33342 stained images to track nuclei of neutrophils migrating under agarose. The x and y position of each cell at each time point is exported as a CSV file. Tracks were filtered and migration parameters were quantified using an in-house MATLAB script. The script first filters cells which travel less than 20 microns as these are mostly broken tracks or background. Additionally, it removes data for cells past 60 min of movie time since the gradient is often dispersed at this stage. The number of cells for a given movie is recorded as the number of tracks after filtering. Migration parameters are then calculated for each track within a movie.

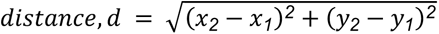

Accumulated distance is calculated as the sum of all distances (*d*) between tracked points using the distance formula.

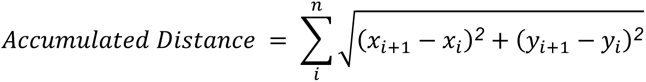

Euclidean distance is calculated by applying the distance formula to the first and last recorded tracked points for a cell.

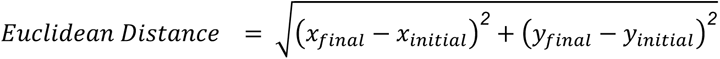

Average speed was calculated by dividing the total distance traveled by the total time elapsed. Time elapsed was calculated as the number of frames times the time interval (30 seconds).

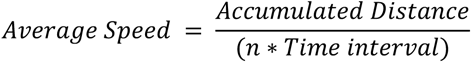

Directionality is calculated as the Euclidean distance divided by the accumulated distance.

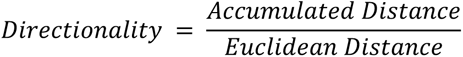

The mass spectrometry data was sorted based on >2-fold change, statistically significant changes (p >0.05) in the LMNA proteome relative to the SCR proteome. The data were further analyzed for the abundance of proteins based on biological functions using the ShineyGO 0.73 gene enrichment analysis platform and were plotted in Fig. S5a-b. Relative protein abundance as observed in Westen blot images was quantified using the Analyze>Gel feature of Image J (Fig. 2e-f, 4a-d, and 5b-c).

All the data presented in this manuscript are derived from a minimum of three independent biological replicates unless otherwise stated. We have employed suitable statistical tests to assess significance, ascertain confidence levels, and evaluate variability within the data sets, with specific details provided in the relevant figure legend.

