## Supplemental Information for "Neutrophils secrete exosome-associated DNA to resolve sterile acute inflammation"

### SUPPLEMENTARY INFORMATION

#### SUPPLEMENTAL FIGURE LEGENDS

**Figure S1**

**Chromatin-like bead-on-string structures are present within MVBS containing ILVs in activated PMNs.**

**a-b.** Transmission electron microscopy images of PMNs migrating towards $LTB_4$ showing the presence of chromatin-like structures in vesicles close to the NE and in cytoplasmic MVBs alongside ILVs. The scale is 5 µm, and it is 200 nm in the inset.

**Figure S2**

**Unlike PMA-induced suicidal NET release $LTB_4$-induced DNA secretion is non-lytic and is PAD4- and NOX2-independent.**

**a-b.** Time-lapse snapshots of human PMNs stained with Mitotracker red CMXros (red, mitochondria), and Hoechst (blue, nuclei) either migrating towards an increasing gradient of $LTB_4$ (**a**) or treated with 100 nM PMA (**b**), in the presence of the membrane-impermeable DNA-binding dye SYTOXgreen (green). Zoomed insets from panel b, show the outline of the cell (red), nuclei (blue), and NETs (green); obtained using CellProfiler-based object segmentation. The scale is 5 µm. See the associated Movies 3 and 5. N=2.

**c.** Graph showing the change in the mean nuclei area extent (nuclear swelling and rupture) over time in PMNs treated with either PMA or migrating towards $LTB_4$. Each dot represents the average value obtained from all the PMNs within 40,000 $µm^2$. Blue and red dots indicate the percentage of PMNs with either no SYTOXgreen staining in the cytoplasm or positive for SYTOXgreen (suicidal NET release).

**d.** Graph showing the percentage of PMNs lysed over time and quantified using CellProfiler-based object segmentation. Datapoints are plotted as mean (red/blue dots) ± s.e.m. (black lines). N=3.

**e.** Graph showing the percent of maximum LDH activity in the culture supernatants of PMNs treated with either DMSO, PMA (20 nM), or $LTB_4$ (100 nM), for 2 hrs. Datapoints from the same experiment are similarly color-coded and are plotted as mean ± s.e.m. and P values calculated using RM one-way ANOVA are shown. N=3.

**f.** Graph showing the changes in the Mitotracker Red intensity in migrating PMNs before, during, and after DNA secretion within 1 hr as shown in panel a. The plotted thick black lines show mean ± s.e.m., with red circles representing 30 datapoints for PMNs migrating towards $LTB_4$. P values calculated using ordinary one-way ANOVA are plotted. N=2.

**g.** Scatter dot plot showing the percent of DNA-secreting PMNs among those migrating towards $LTB_4$ within an observation window of 40,000 $µm^2$ for 1 hr in the presence or absence of PAD4 and NOX2 inhibitors, GSK484 (2 µM) and GSK2795039 (10 µM) respectively. Datapoints from the same experiments are similarly color-coded and are plotted as mean ± s.e.m. and P values calculated using RM one-way ANOVA are shown. N=3. See associated Movie 6.

**h.** Scatter dot plot showing the percent of DNA-secreting cells among those migrating towards $LTB_4$ within an observation window of 40,000 $µm^2$ for 1 hour in the presence or absence of various cell death pathway inhibitors namely, Z-DEVD-FMK (apoptosis, 10 µM), ferrostatin 1 (ferroptosis, 5 µM), and GSK872 (necroptosis, 10 µM) as indicated on the x-axis. Datapoints from the same experiments are similarly color-coded and are plotted as mean ± s.e.m. and P values calculated using mixed effect analysis are shown. N≥3. See associated Movie 7.

**Figure S3**

**No change in cellular and nuclear morphology during and after SEADing.**

**a-b.** Scatter dot plots showing the change in the nuclear form factor (**a**) and cell eccentricity (**b**) during DNA secretion compared to PMNs without DNA secretion. The red circles indicating 45 datapoints for DNA-secreting cells and 31 data points for cells without DNA secretion are plotted as mean ± s.e.m., with black lines representing error bars. P values obtained using ordinary one-way ANOVA for DNA-secreting cells and the Mann-Whitney test in DNA-secreting before condition with no DNA secretion are plotted on the graph. N=5.

**Figure S4**
***LMNA* KO dHL60 cells have a nuclear morphology closer to PMNs relative to SCR dHL60 cells.**
**a.** Representative western blot images showing the levels of lamin A/C, lamin B1, lamin B2, and LBR in cell lysates obtained from SCR, *LMNA* KO, and *LMNA/LBR* KO dHL60 cells. GAPDH is used as a loading control. The molecular weight in kilodaltons (kDa) is located on the left side of the panels.
**b.** Representative Airyscan microscopy images of PMNs, SCR, *LMNA* KO, and *LMNA/LBR* KO dHL60 cells migrating towards fMLF, fixed and immunostained for LBR (magenta) and co-stained with Hoechst (cyan, nucleus). Images are presented as a "sum of slices" projection of the acquired z-stack. Dotted white/black outlines indicate cell shape. The scale is 5 µm.
**c-f.** Scatter dot plots showing the changes in the NE to cytoplasm LBR intensity ratio (**c**), nuclei form factor (**d**), NE invaginations (**e**), and heterochromatin spots (**f**), in dHL60 neutrophils as shown in panels a-b. The red circles indicate a minimum of 29 datapoints for PMN, 27 for SCR, 41 for *LMNA* KO, and 15 datapoints for *LMNA/LBR* KO dHL60 cells plotted as mean ± s.e.m. P values calculated using ordinary one-way ANOVA are shown. N=3.

**Figure S5**
***LMNA* KO dHL60 cells exhibit a proteome profile closer to PMNs and migrate better relative to SCR dHL60 cells.**
**a-b.** Graph showing the fold change in the gene ontology (GO, biological process) profile of the associated downregulated (**a**) and upregulated (**b**) proteins in *LMNA* KO dHL60 cells relative to SCR dHL60 cells, quantified using tandem mass tagging (TMT)-based mass spectroscopy analysis.
**c-f.** Scatter dot plots showing the changes in the number of cells migrating towards fMLF for 1 hr (**c**), median speed (**d**), median directionality (**e**), and cell eccentricity (**f**). Data is plotted as mean ± s.e.m. of 9, 8, and 5 datapoints (red circles) for SCR, *LMNA* KO, and LMNA*/LBR* KO samples. N≥5. For cell eccentricity, a total of 29, 45, and 16 datapoints for SCR, *LMNA* KO, and *LMNA/LBR* KO, were pooled from 3 independent experiments. The P values calculated using mixed-effect analysis (c-e) and ordinary one-way ANOVA (f) are shown.

**Figure S6**
**DNase I-induced disruption of secreted DNA impedes monocyte infiltration during the resolving phase of TPA-induced ear inflammation.**
**a-b.** Representative Airyscan microscopy images of acetone/TPA-treated mice ear cryosections (20 µm thick) treated for the indicated duration in mice injected with either DNase I or PBS, showing the presence and relative localization of (**a**) citrullinated histone (magenta, secreted DNA) and (**b**) F4/80 (magenta, macrophages) with DAPI (gray, nuclei). Images are presented as a 'sum of slices' projection of acquired z-stacks. The scale is 100 µm. N=3.

**Figure S7**
**PLA of FLAP-positive extracellular structures with dsDNA in TPA-treated ears.**
**a.** Schematic illustrating the antibody binding sites on FLAP and the principle of the PLA used to assess the presence of SEADs in inflamed ears.
**b.** Representative Airyscan microscopy images of 20 µm thick cryosections of mouse ear treated with TPA for 12 hrs, processed for PLA, and showing the status of PLA dots (magenta) in either

FLAP- or dsDNA-only antibody conditions, used to test the non-specific PLA signal. Sections were co-stained with DAPI (nucleus). The scale is 100 µm. N=3.

### MOVIE LEGENDS

**Movie 1**
**Tomogram and segmentation of data shown in Fig. 2c-d.**

**Movie 2**
**Non-lytic, repetitive, and rapid secretion of DNA from chemotaxing PMNs.**
PMNs stained with CellMask orange (PM) and Hoechst 33342 (nuclei) migrating towards $LTB_4$ in the presence of SYTOXgreen imaged using Airyscan microscopy. Images acquired at 30-sec intervals are presented as 3 frames per second. Scale 5 µm.

**Movie 3**
**The diverse phenotypes of DNA secreted from chemotaxing PMNs.**
PMNs stained with Mitotracker Red CMXros (mitochondria) and Hoechst 33342 (nuclei) migrating towards $LTB_4$ in the presence of SYTOXgreen imaged using confocal microscopy. Images acquired at 30-sec intervals are presented as 3 frames per second. The scale is 5 µm. Cyan arrow marks DNA-trails, white triangle marks 'DNA-blobs', and hollow white triangles mark 'attached DNA-blobs'.

**Movie 4**
**The secretion of DNA from chemotaxing PMNs is dependent onSMase activity.**
PMNs stained with CellMask orange (PM) and Hoechst 33342 (nuclei) migrating towards $LTB_4$ in the presence of SYTOXgreen, and either DMSO (vehicle control) or GW4869 (nSMase inhibitor) imaged using confocal microscopy. Images acquired at 30-sec intervals are presented as 3 frames per second. Scale 5 µm.

**Movie 5**
**PMA-induced suicidal NET release in PMNs.**
PMNs stained with CellMask orange (PM) and Hoechst 33342 (nuclei) stimulated with PMA (100 nM) in the presence of SYTOXgreen and imaged using Airyscan microscopy. Images acquired at 18-sec intervals are presented as 3 frames per second. Scale 5 µm.

**Movie 6**
**The secretion of DNA from chemotaxing PMNs is independent of PAD4 and NOX2 activity.**
PMNs stained with CellTrackerCMPTX (cell) and Hoechst 33342 (nuclei) migrating towards $LTB_4$ in the presence of SYTOXgreen, and either DMSO, PAD4 inhibitor and NOX2 inhibitor imaged using confocal microscopy. Images acquired at 45-second intervals are presented as 1 frame per second.

**Movie 7**
**Effect of various cell death pathway inhibitors on the secretion of DNA from chemotaxing PMNs.**
PMNs stained with CellMask orange (PM) and Hoechst 33342 (nuclei) migrating towards $LTB_4$ in the presence of SYTOXgreen, and either DMSO, apoptosis, ferroptosis, or necroptosis inhibitors

imaged using confocal microscopy. Images acquired at 30-sec intervals are presented as 3 frames per second.

**Movie 8**

**LBR loss inhibits DNA secretion in chemotaxing dHL60 cells.**

SCR, *LMNA* KO, and *LMNA/LBR* KO dHL60 cells stained with CellMask orange (PM) and Hoechst 33342 (nuclei) migrating towards $LTB_4$ in the presence of SYTOXgreen imaged using confocal microscopy. Images acquired at 30-sec intervals are presented as 3 frames per second.

**Movie 9**

**Histone acetylation mediates the secretion of DNA from chemotaxing PMNs.**

PMNs stained with CellMask orange (PM) and Hoechst 33342 (nuclei) migrating towards $LTB_4$ in the presence of SYTOXgreen, and either DMSO, HAT, or HDAC inhibitors imaged using confocal microscopy. Images acquired at 30-sec intervals are presented as 3 frames per second.

**Movie 10**

**Effects of MK886 and DNase I treatment on PMN chemotaxis.**

PMNs stained with Hoechst 33342 migrating towards $LTB_4$ in the presence of either DMSO, DNase I, MK886, or MK886 + DNase I, were imaged using fluorescence microscopy and analyzed using Trackmate on ImageJ. Images acquired every 45 sec are presented at a rate of 5 frames per second. Circles indicate the individual cells as identified by Trackmate, and lines are color-coded for the duration of migration from blue (earlier) to red (later). The scale is 200 μm.
