## Supplementary figures and images for "Neutrophils secrete exosome-associated DNA to resolve sterile acute inflammation"

### Supplemental Figures

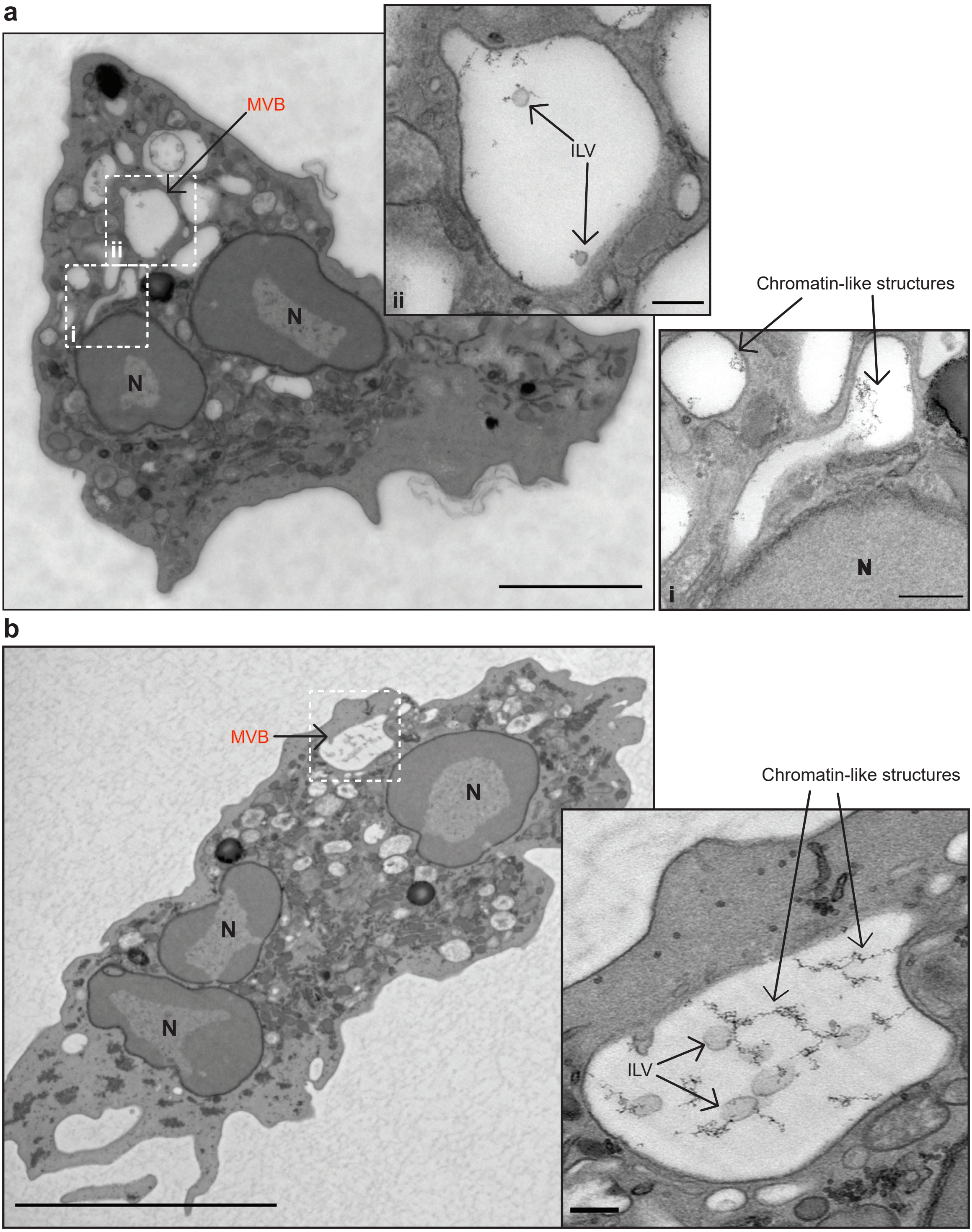

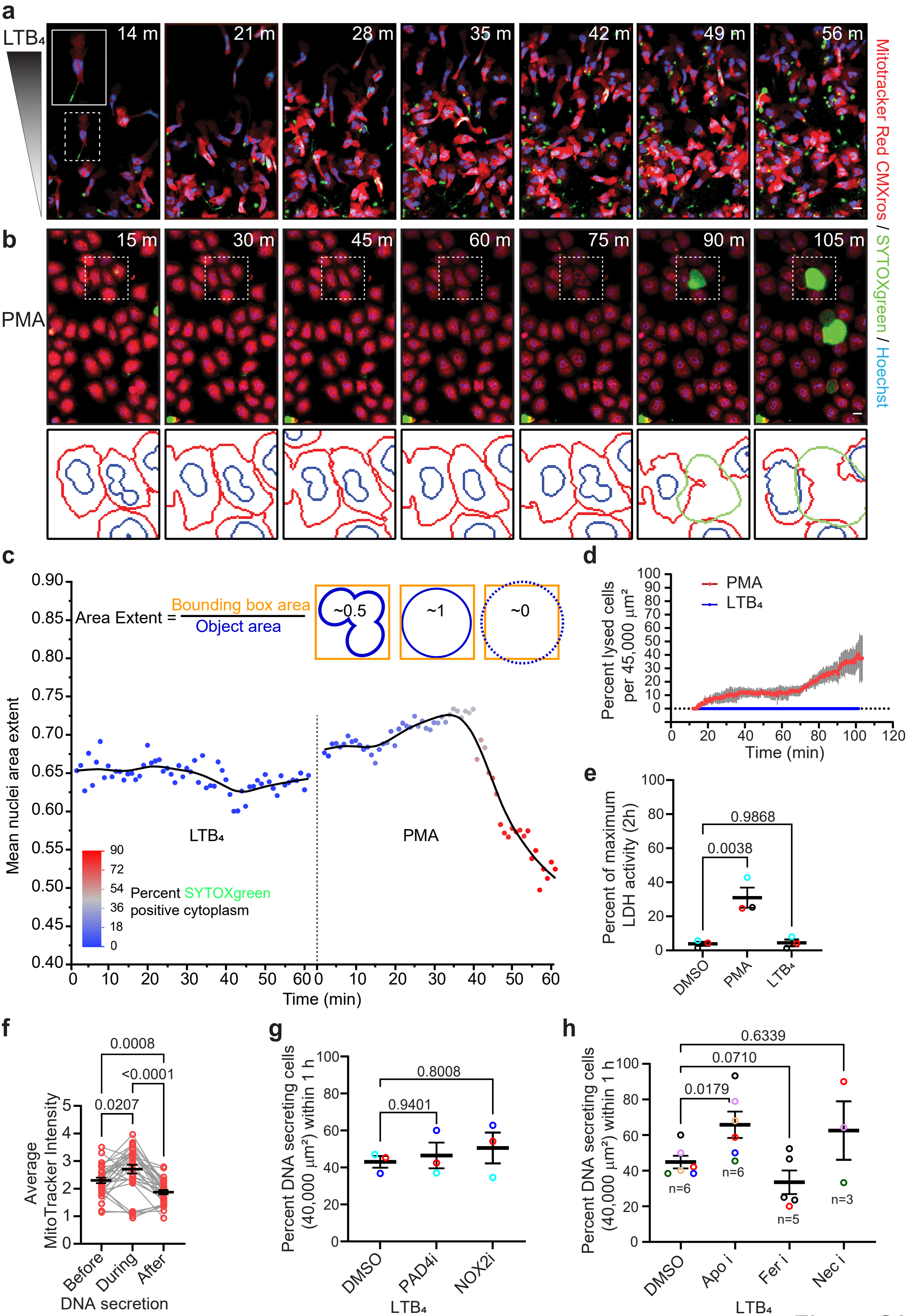

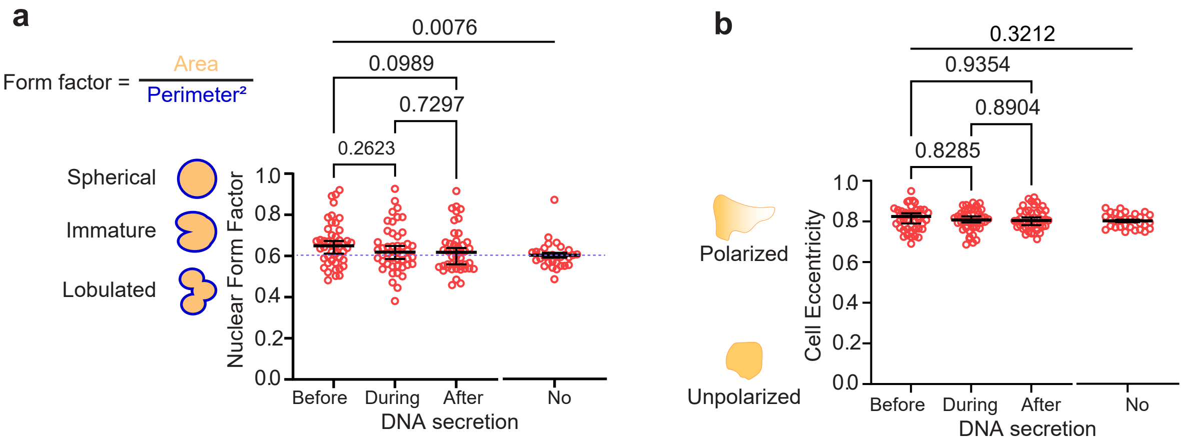

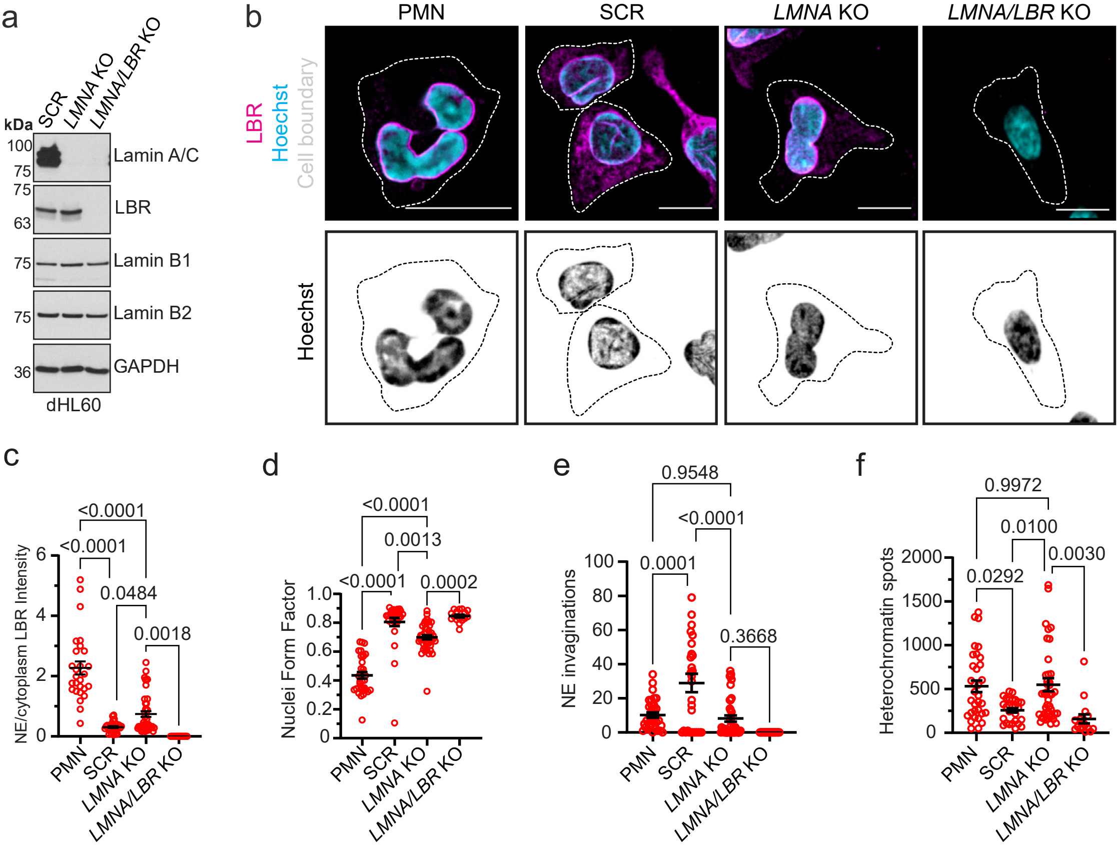

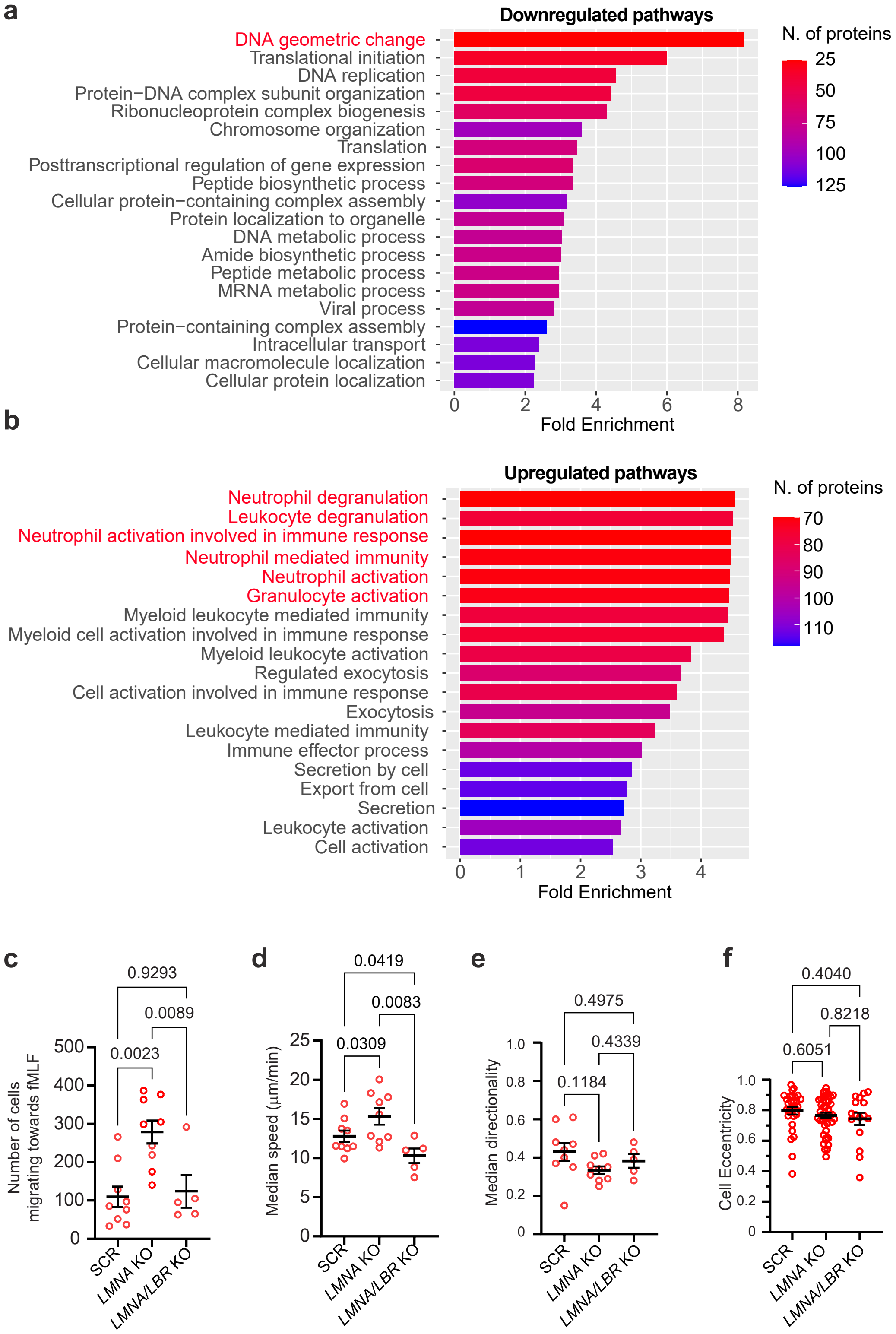

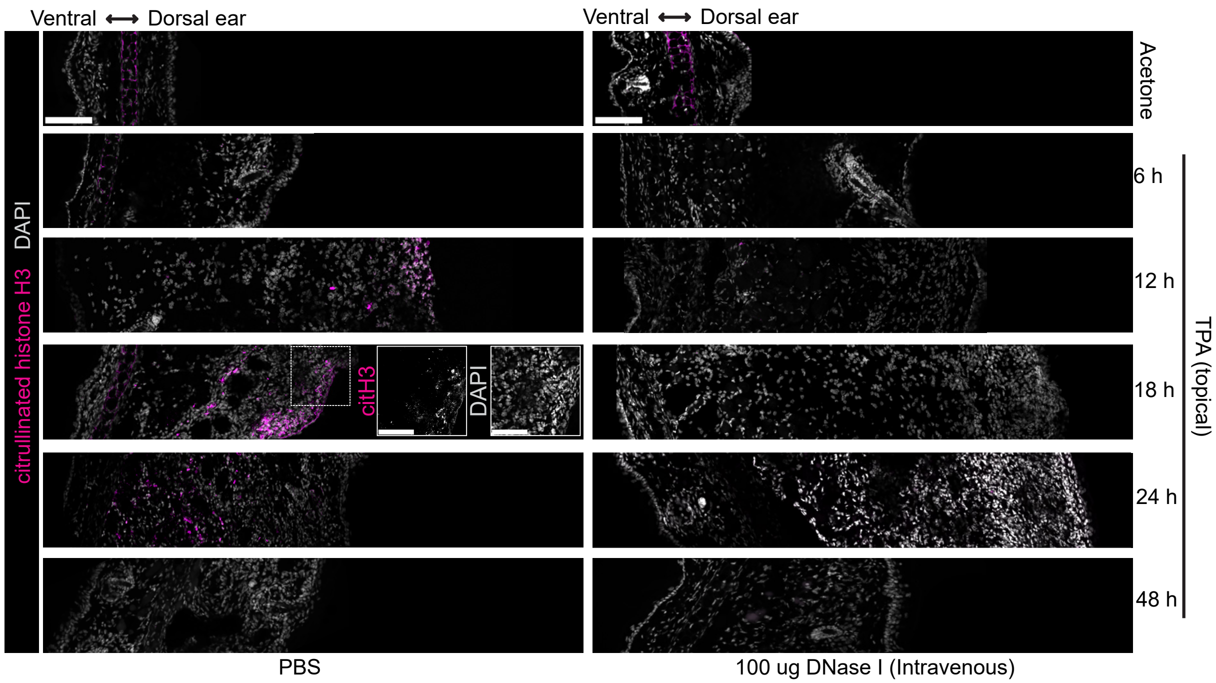

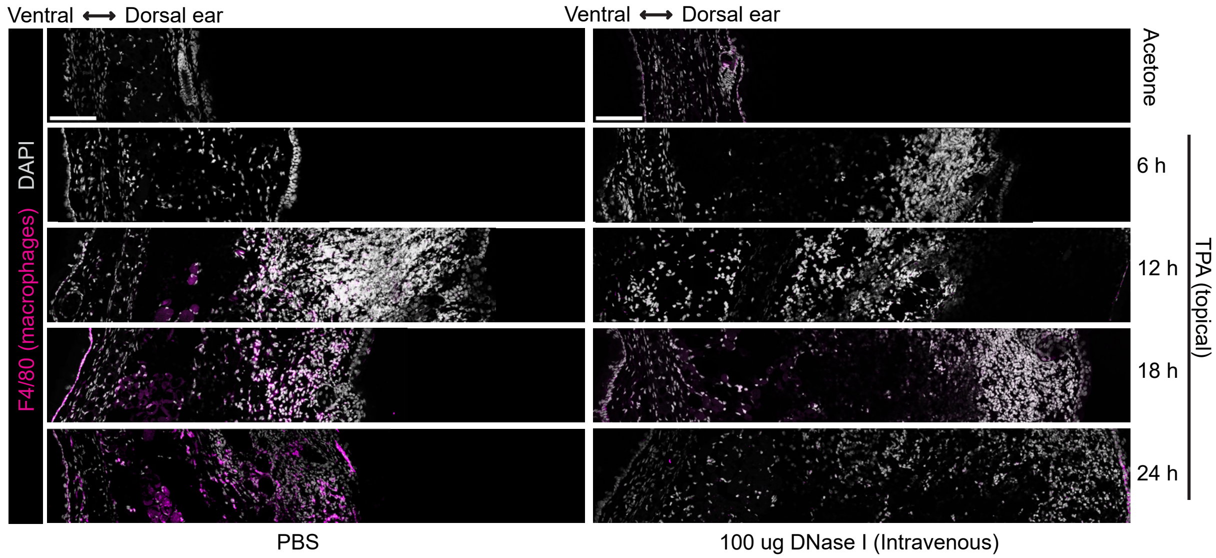

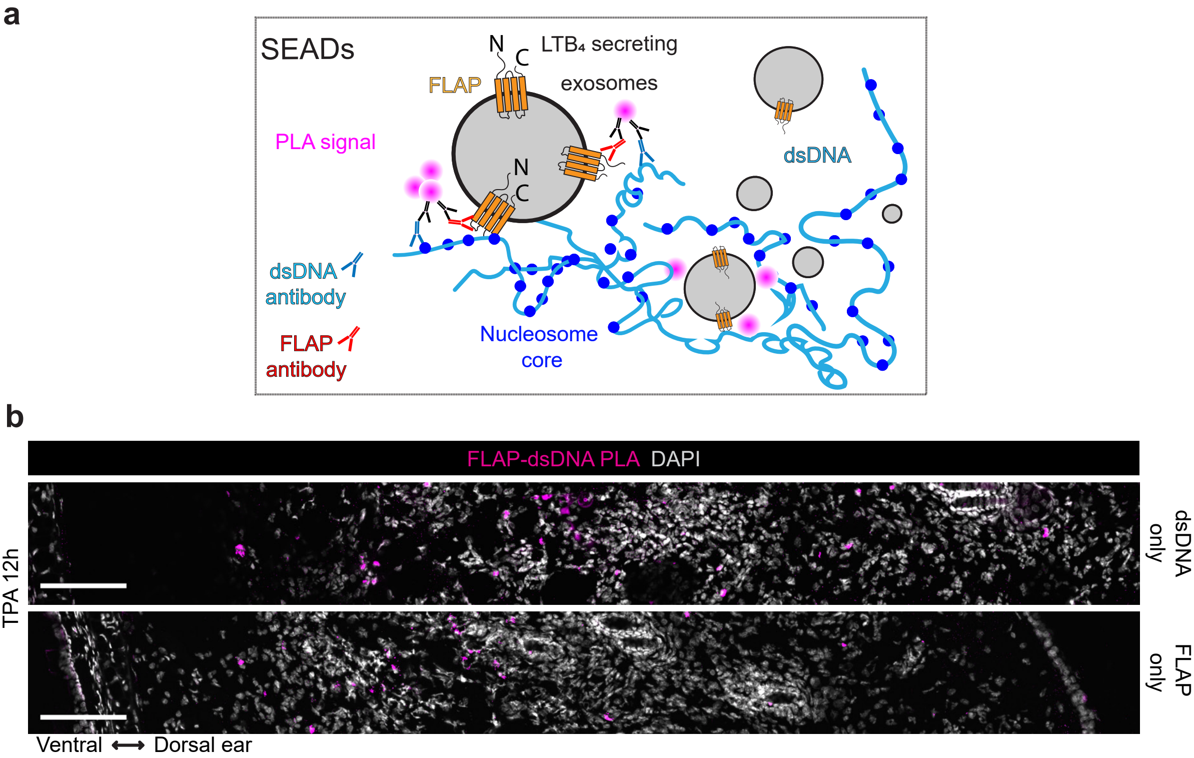
